# A weighted multi-trait approach for heterotic grouping of maize inbred lines under *Striga* infestation and optimum environments

**DOI:** 10.64898/2026.05.15.725596

**Authors:** Adamu Masari Abubakar, Idris Ishola Adejumobi, Wende Mengesha, Silvestro Meseka, Muhyideen Oyekunle, Shehu Garki Ado, Tégawendé Odette Bonkoungou, Baffour Badu-Apraku, John Derera

**Affiliations:** International Institute of Tropical Agriculture (IITA), PMB 5320, Oyo Road, Ibadan, Oyo State, Nigeria; Department of Plant Science, Institute for Agricultural Research, Ahmadu Bello University, P.M.B. 104, Zaria, Kaduna State, Nigeria; Institut de l’Environnement et de Recherches Agricoles (INERA), 01 B.P.910, Bobo-Dioulasso 01, Bobo-Dioulasso, Burkina Faso

**Author notes:** Corresponding author (W.M.). (A.M.A.); (I.I.A.); (S.M.); (B.B.A.); (J.D.) (A.M.A.); (M.O.); (S.G.A.) (T.O.B.).

**Keywords:** Combining ability, correlation, gene action, heterotic group, *Striga*-resistant

## Abstract

Maximum utilization of existing genetic variability in a breeding program depends on the efficient classification of the inbred lines into heterotic groups, particularly under stress conditions. This study applied practical breeding approaches to determine the mode of genetic inheritance for *Striga* resistance and proposes a weighted heterotic grouping method based on the general combining ability of multiple traits (WHGCAMT) and compares its effectiveness with other existing methods in classifying the inbred lines into heterotic groups in *Striga*-infested and optimum environments. Using Diallel design IV, 300 crosses were generated from 21 inbred lines and 4 standard testers. The crosses, along with six checks, were evaluated in an 18 × 17 alpha lattice design with two replications at two locations, in both artificial *Striga*-infested and *Striga*-free environments. The inbred lines were genotyped using DArTtag SNP markers. Phenotypic and genotypic data were analyzed using R. Analysis of variance revealed significant mean squares for hybrid, general combining ability (GCA), specific combining ability (SCA) and their interactions with environment. Significant positive and negative GCA and SCA effects were detected for grain yield and other measured traits. However, a larger proportion of additive gene action than non-additive gene action was observed for grain yield and most measured traits. The analysis of molecular variance also showed substantial genetic differences within and between clusters. Except for HSCA, the mean grain yield between the inter-group and intra-group hybrids was significant for each method. Pairwise comparison of the inter- and intra-group hybrids of all the methods showed significant differences between the WHGCAMT and all other methods in most cases. WHGCAMT consistently produced higher-yielding inter-group hybrids and lower-yielding intra-group hybrids, achieving breeding efficiency improvements of 55.8%, 4.3%, 15.7%, and 11.4% over the HSCA, HSGCA, HGCAMT and molecular marker methods, respectively, under *Striga* infestation. Thus, WHGCAMT offers more precise, reliable and biologically meaningful heterotic groups among early-maturing maize inbred lines.

## Introduction

Maize (*Zea mays* L.) hybrid production has the potential to address the challenges of the world’s rapidly growing population. The production of maize ranks first in Africa and Latin America while in Asia it ranked third after rice and wheat (1). In sub-Saharan Africa (SSA), maize is contributing to the food security and livelihoods of millions of people (2). However, persistent challenges of *Striga hermonthica* (purple witchweed) steadily decreases its production in the savannas of SSA (3). The obligate hemiparasitic plant, *S*. *hermonthica,* parasitizes the root systems of cereals, resulting in a notable decrease in productivity. The parasitic weed can cause severe yield losses ranging from 10 to 100%, particularly when infestation is severe during the vegetative growth stage and farmers have reportedly abandoned fields due to *Striga* attack (4,5). The use of improved hybrids with enhanced resistance to *S. hermonthica* has been demonstrated as the most sustainable approach for minimizing yield reductions on farmer’s field with limited resources (6). This led to a continuous need for genetic improvement of various economically important traits, particularly under hemiparasitic attack, to sustain hybrid maize production.

Sustainable hybrid maize production depends heavily on the efficient classification of the parental lines into heterotic groups (HGs) (7,8). Heterotic grouping enhances the likelihood of combining genetically diverse but complementary parents, leading to superior hybrid performance in yield, agronomic traits and stress tolerance (9–11). It is therefore essential to have knowledge of the pattern of HGs among inbred lines in a breeding program for selecting parents to develop heterotic hybrids, identifying sources of useful alleles for introgressive hybridization and choosing suitable inbred or hybrid testers (12–14). Inbred lines can be classified into distinct HG by either molecular based marker or phenotypic traits (15). Molecular-based approaches classify lines based on genetic distance derived from the molecular markers (16), which is very useful for describing heterotic groups and studying relationship among inbred lines at the molecular level. The phenotypic approach classifies of inbred lines using BLUP estimates called combining ability. Several researchers have reported different methods. Vasal et al., (14) proposed heterotic grouping based on specific combining ability (SCA) effects of grain yield (HSCA) to classify inbred lines into HGs. Several researchers (13,17,18) have used this method, efficiently classified inbred lines into HGs. However, the method has often been found to be influenced by the interaction between the two parents and between hybrids and environment, which often leads to the placement of the same inbred lines into different heterotic groups (7,11,19). To address the inadequacies of the HSCA approach, Fan et al., (11) proposed the heterotic grouping based on specific and general combining abilities for grain yield (HSGCA) method. The HSGCA identified the best parental combinations, leading to the development of high-yielding and resilient hybrids (11). This method has been shown to be more effective than the HSCA method (10,11,15,19) or molecular SSR (19) and SNP (20) markers for classifying inbred lines into distinct heterotic groups. The limitation of heterotic grouping of the inbred lines using HSCA and HSGCA methods is the involvement of single traits (primarily grain yield) in its calculation (21). Grain yield is a complex trait controlled by polygenes and has low heritability, especially under stress environments (4,12,21,22). Using grain yield alone, improvement would be difficult through direct selection, particularly under drought and *Striga*-infested conditions. Therefore, it is more effective to use component traits that have a strong correlation with grain yield for genetic improvement (21). It has been documented that by including other secondary traits, a more precise classification of the inbred lines into heterotic groups is achieved (21). In this regards, Badu-Apraku et al. (21) proposed heterotic grouping based on general combining ability of multiple traits (HGCAMT). The HGCAMT method is a specialized approach developed to quantify the genetic relationships among genotypes by utilizing standardized additive genetic effects across multiple traits. This method assigns genotypes into their distinct HGs by computing genotype-specific effects using the HGCAMT’s formula and subsequently applying multivariate clustering analysis, such as hierarchical or k-means clustering. Despite its simplicity and utility, the HGCAMT method has some limitations and needs to be address.

HGCAMT implicitly assigns equal weight to all traits, irrespective of their relative biological, agronomic and economic importance and it treats all deviations from the mean equivalently. However, not all traits contribute equally to breeding progress, particularly under stress conditions (23,24). In maize breeding, for example, grain yield is the primary target trait, yet it is strongly influenced by several secondary traits such as number of ears per plant, *Striga* damage ratings, and stay-green (25,26). Consequently, assigning equal importance to all traits may hot fully ignored breeder priorities or selective objectives. Incorporating trait-specific weights allows breeders to emphasize traits with greater economic or adaptive value while retaining complementary information from secondary traits. This flexibility ensures that heterotic grouping reflects both underlying genetic divergence and practical breeding priorities, thereby increasing the likelihood that inter-group hybrids will combine high yield potential with resilience to biotic and abiotic stresses (27).

Additionally, the HGCAMT approach often utilizes distance matrices that do not account for inter-trait correlations, which can distort the distance matrix and lead to less reliable grouping outcomes, particularly in the presence of multicollinearity (28–30). In multi-trait heterotic grouping, several measured traits are often strongly correlated (31,32). For example, grain yield had a strong positive correlation with ears per plant and a negative correlation with *Striga* damage ratings (25,33). Although these correlations are biologically meaningful, failure to account for them in clustering analyses can lead to statistical redundancy, in which correlated traits exert disproportionate influence on distance calculations (34). Such redundancy may obscure true genetic divergence among inbred lines and introduce biased heterotic grouping outcomes (34). Incorporating the inverse covariance matrix into the distance matrix allows trait correlation to be explicitly accounted for such that correlation dimensions are down-weighted while independent dimensions retain their full influence (35–37). This approach ensures each trait contributes unique information to the grouping process, resulting in balanced, reliable and more precise classification. Therefore, the objectives of this study were to: (i) determine the mode of inheritance for grain yield and other *Striga*-adaptive traits in 25 early maturing inbred lines; (ii) propose a weighted HGCAMT approach and compare its effectiveness with other existing methods in classifying inbred lines into heterotic groups in *Striga*-infested and optimum environments.

## Materials and Methods

### Plant materials

The genetic materials comprised 25 inbred lines involving four standard testers (TZdEI 352, TZEI 18, TZEI 31 and TZEI 7), two newly identified testers (TZEI 2250 and TZEI 2238) and 19 newly developed inbred lines. The testers varied significantly in their reaction to *Striga*. TZdEI 352, TZEI 18 and TZEI 2250 are *Striga-*resistant testers, while TZEI 7 and TZEI 31 are good combiners under low soil nitrogen (low N) and optimum environments but susceptible to *Striga* parasitism (25). The tester TZEI 2238 was recently identified as *Striga* susceptible tester (unpublished). In addition to the standard testers, the other 21 lines were developed using the pedigree method through repeated selfing aided by marker-assisted selection (MAS) and field-testing. At S6, the inbred lines were screened under artificial *Striga* infestation at two locations. Based on the result, TZEI 2250, TZEI 2238 and the other 19 lines varied significantly in grain yield and other *Striga*-adaptive traits. They were advanced to S_7_ and utilized in this study. The detailed description of all the 25 inbred lines is presented in (Table S1).

### Experimental sites

The study was carried out at Mokwa (9^◦^210 N and 5^◦^1 0 E, altitude 188 m.a.s.l.) in 2022 and Abuja (9^◦^150 N and 7^◦^200 E, altitude 431 m.a.s.l.) in 2023, Nigeria. The average annual rainfall in Abuja and Mokwa was 1,513 mm and 1,200 mm, respectively. The soil at Abuja was sandy, while the soil at Mokwa was sandy loam. The detailed characteristics of the study sites were described by (38)

### Cross-generation and evaluation

The 25 inbred lines were crossed in Diallel fashion using method IV to develop 300 single cross hybrids. The 300 crosses plus 6 checks were planted in 18 × 17 alpha lattice design with two replications. The trials were conducted at Abuja and Mokwa in Artificial *Striga*-infested and *Striga*-free environments, which were located 200 m apart. In each trial, the hybrids were planted in a 3 m-long single-row plot with an inter-row spacing of 0.75 m and an intra-row spacing of 0.40 m. The *Striga*-free plots were treated with ethylene gas two weeks before planting to eliminate (if available), any potential *S. hermonthica* seeds present in the soil. The seeds of *S. hermonthica* used for infestation in each year were collected in the previous year from farmers’ sorghum fields from each of the locations. Infestation was carried out by applying 8.5 g of sand-mixed *S. hermonthica* seed inoculum to holes roughly 6 cm deep, 10 cm wide, and 0.40 m apart. The estimated number of germinable *Striga* seeds per hill was 3000 (26). Three seeds of each hybrid were placed into a hole infested with the sand mixed with *S. hermonthica* seeds and covered with soil. Plots were thinned to two plants per hill two weeks after sowing to attain a population density of 66,666 plants per ha. For fertilizer application, the *S. hermonthica* field received a split dosage application of NPK 15-15-15 with the first dose application done at three weeks after planting (WAP) and second application at five WAP. Both applications provided the plants with 60 kg/ha N, 60 kg/ha P and 60 kg/ha K. The Striga free field received normal fertilizer dose of 60 kg/ha of N, P, and K respectively at two WAP with another 60 Kg/ha of N applied as topdress with Urea 46-0-0. Weeds other than *S. hermonthica* were manually removed throughout the cropping seasons from the *S. hermonthica* field while herbicide (contact) was used for weed control in *Striga*-free plots at the rate of 10 ml/ 20-liter knapsack tank in the *Striga*-free field.

### Data collection

Data were collected in both *Striga*-infested and *Striga*-free plots on: plant height (PHT), ear height (ETH), number of emerged *Striga* plants (ESP), *Striga* damage symptom (SDS), plant aspect (PASP), ear aspect (EASP), husk cover (HCV), ears per plant (EPP), anthesis silking interval (ASI) and grain yield (GYD). PHT was measured in cm from five randomly selected plants in a plot as the distance from the base of the plant to the height of the first tassel branch. Ear Height was measured in cm from five randomly selected plants in a plot as the distance from the base of the plant to the node bearing the upper ear. Ear Aspect was recorded as number of emerged *S. hermonthica* plants in each infested plot at 10 weeks after planting (WAP). *Striga* damage symptom was visually rated on a per-plot basis in each infested plot at 10 WAP using a scale of 1 to 9, where 1 = normal plant growth with no visible host plant damage symptom, and 9 = all leaves completely scorched, resulting in premature death. PASP was rated on per - plot basis on a scale of 1 to 9, where 1 = excellent plant with large and uniform ears, low ear placement, shorter plants, resistant to foliar diseases and little or no stalk and root lodging, and 9 = plants with small and variable ears. EASP was rated on per-plot basis on a 1 to 9 scale, where 1 = clean, uniform and large ears, and 9 = rotten, variable and small ears. HCV is rated on per-plot basis on a scale of 1 to 9, where 1 = husks tightly arranged and extended beyond the ear tip, and 9 = loose husks with ear tips exposed. EPP was calculated by dividing the number of harvested ears by the number of plants harvested per plot. ASI was calculated as the absolute difference between days to 50% silking and days to 50% anthesis. GYD was computed in kg/ha from the weight of shelled grain adjusted to 80% shelling percentage and corrected for 15% moisture content (25)

### DNA extraction and genotyping

Young and fully expanded leaves were sampled from the plot of each inbred in the nursery using the bulk method at 3 weeks after planting and stored at – 80 °C for 72 hours before lyophilization. The freeze-drying (lyophilization) was performed using a FreeZone Freeze Dryer (Labconco, USA). Freeze-dried samples were shipped to Diversity Array Technology Ltd, Australia, for genotyping. The genotyping was performed using DArTag technology, a reduced-representation sequencing method that reduces genome complexity using restriction enzyme digestion, followed by next-generation sequencing. Sequencing libraries were prepared using a proprietary combination of restriction enzymes to target low-copy genomic regions, and the libraries were sequenced on an Illumina platform. Both SNP and SilicoDArT (presence/absence) markers were identified using DArT’s proprietary bioinformatics pipeline, as described by Kilian et al. (39).

### Data analysis and heterotic grouping

#### a. Analysis of molecular variance (AMOVA) and heterotic grouping using molecular markers

In total, 3,305 SNP markers in a HapMap-format file received from the DArT platform were converted to the Variant Call Format (VCF) for downstream analyses. Quality control filtering was applied to retain only high-quality balletic SNPs. Markers with sequence depth < 5 (based on DArT quality metrics), missing data > 20%, minor allele frequency (MAF) < 5%, were excluded. Indel markers were also removed. After filtering, 1696 high-quality SNPs distributed across the 10 chromosomes were retained for further analyses. Summary statistics such MAF, observed and expected heterozygosity (Ho and He), for both SNP markers and genotypes were assessed using”– freq” and “–hardy” functions implemented in Plink v2.0 (40). The filtered SNP dataset was subsequently converted to numeric format using GAPIT package (41) in R.

#### b. Analysis of variance (ANOVA), specific and general combining abilities

After quality checks, the phenotypic data were subjected to analysis of variance (ANOVA) across *Striga*-infested and optimal environments using a mixed model procedure. The environment, replicate and block were considered as random factors, whereas the 300 single-cross hybrids as fixed. In this analysis, the sources of variation were partitioned into the environment, genotype, additive, dominance and their interactions with the environment. The model used for the analysis is presented in equation 1.

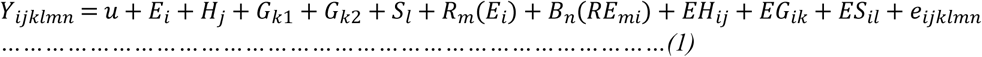

Where:

𝑌_𝑖𝑗𝑘𝑙𝑚𝑛_: Phenotypic observation.

𝑢: Grand mean.

𝐸_𝑖_: Random effect of ith environment.

𝐻_𝑗_: Fixed effect of jth hybrid (genotype).

𝐺_𝑘1_𝑜𝑟 𝐺_𝑘2_: GCA effect of k1th or k2th parent.

𝑆_𝑙_: SCA effect for the cross between the k1th and k2th parents.

𝑅_𝑚_(𝐸_𝑖_): Random effect of the replicate nested within environment.

𝐵_𝑛_(𝑅𝐸_𝑚𝑖_): Random effect of the block nested within replicate and environment.

𝐸𝐻_𝑖𝑗_: Interaction between genotype and environment.

𝐸𝐺_𝑖𝑘_: Interaction between GCA and environment.

𝐸𝑆_𝑖𝑙_: Interaction between SCA and environment.

𝑒_𝑖𝑗𝑘𝑙𝑚𝑛_: Residual error.

The mean squares for each source of variation across *Striga*-infested and optimum environments were estimated for the 300 Diallel crosses. The general combining ability (GCA) of the parents and specific combining ability (SCA) of the crosses were estimated based on Griffing’s method 4 mixed B (42). The analysis was performed for grain yield and all other traits following the Henderson method using the Analysis of Genetic Designs with R (AGD-R) for Windows software version 5 (43). Thereafter, the best linear unbiased estimations (BLUEs) of the 300 hybrids and 6 checks were extracted using the Multi-Environment Trials Analysis in R (META-R) v2.1 R package software (44).

#### c. Contribution of GCA and SCA

The relative importance of GCA and SCA was calculated using the equation:

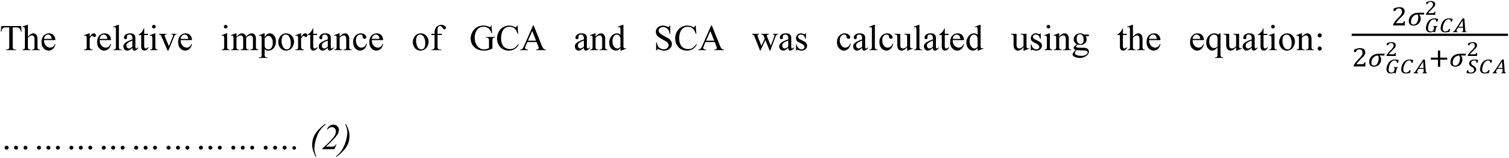

Modified from (45) by (46), where 2σ^2^ is the variance effects derived from the mean squares of GCA and σ^2^ is the variance effects derived from the mean squares of SCA.

#### d. Heterotic grouping using HSCA

Inbred lines were assigned into heterotic groups based on the SCA effects of their hybrids. According to Vasal et al., (14), a single cross-hybrid with positive SCA effects for grain yield indicated that the parents that constituted the hybrid are in opposite heterotic groups, whereas negative SCA effects for grain yield indicated that the parents are in the same heterotic group.

The two inbred parents constituted a single-cross hybrid showing the highest positive SCA effect for grain yield in this study were designated as reference testers for heterotic groups A (HG A) and B (HG B). The remaining inbred lines were classified based on their SCA effects when crossed with these two testers. An inbred line exhibiting a positive SCA effect with tester A and a negative SCA effect with tester B was assigned to HG B, and vice versa. A third group (HG C) was formed when an inbred line showed similar SCA effects with both testers, either both positive or both negative. In such cases, a t-test was performed to compare the two SCA values. If the difference was significant (*p ≤ 0.05*), the line was assigned to the heterotic group corresponding to the lower (more negative) SCA effect. If the difference was not significant (*p > 0.05*), suggesting no clear affinity to either tester, the line was placed in HG C, representing an intermediate or undefined heterotic group.

#### e. Heterotic grouping using HSGCA

The assignment of inbred lines into heterotic groups using the HSGCA method proposed by Fan et al., (11) was done as follows:

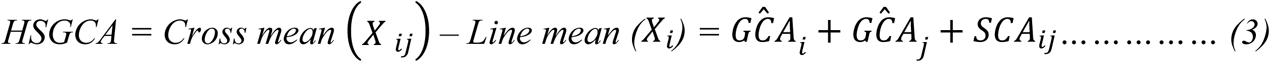

#### f. Heterotic grouping using HGCAMT

The GCA effects of measured traits that showed significant differences (*p<0.05*) for genotypes were standardized and use to estimate HGCAMT as follows:

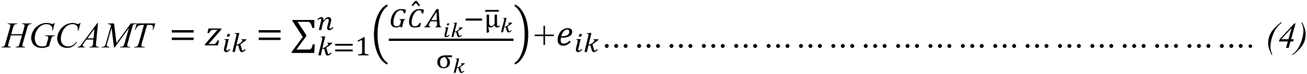

*Where* 𝑧_𝑖𝑘_ *= HGCAMT score for inbred line* i *across all* n *traits,* 𝐺𝐶𝐴_𝑖𝑘_*= GCA effect of inbred* i *for trait* k, μ_𝑘_*= Mean GCA across all inbreds for trait* k, σ_𝑘_ *= Standard deviation of GCA values for trait* k, 𝑒_𝑖𝑘_*= residual (usually small and regularly ignored), ∑ = is over the* n *traits (sum of all the traits)*.

HGCAMT formula above quantify the genetic relationship among inbred lines based on total genetic merit of the GCA across measured traits. However, the HGCAMT assumed equal weight for all the measured traits including grain yield. Better if weight is introduced for the most important traits as:

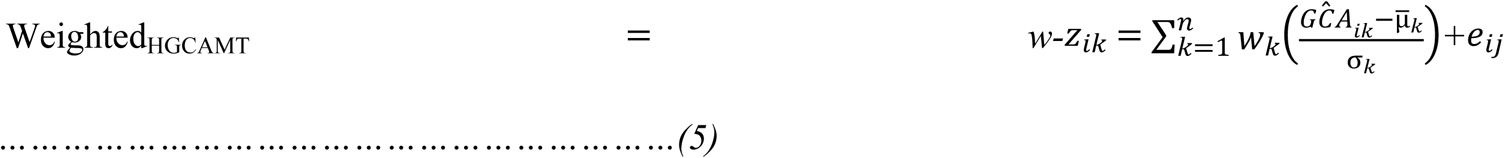

Where 𝑤_𝑘_ *= relative importance or economic weight of the trait k*.

To avoid double-counting correlated traits in the distance computation, a Mahalanobis-type metric in which the difference vector between two inbreds is premultiplied by the inverse of the trait covariance matrix was used. This procedure does not remove correlation from the raw data, but it adjusts the distance measure to account for correlation. In this way, each trait dimension contributes unique information to heterotic grouping, preventing inflated divergence estimates due to trait redundancy.

#### g. Heterotic grouping using WHGCAMT

Heterotic grouping using WHGCAMT emphasizes priority traits and considers trait relationships, ensuring that redundant traits do not dominate the grouping process, as follows:

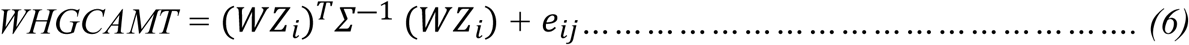

*Where* 𝑊 *= a diagonal matrix of weighted trait* 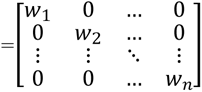

𝑍_𝑖_*=* 𝑧_𝑖_ (𝑧_𝑖1_,…………𝑧_𝑖𝑛_)^⊤^ *is the vector of standardized trait values for genotype i.* 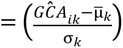 as in HGCAMT adjusting for differences in scale measurement of the traits

𝛴^―1^ *= the inverse covariance matrix of the standardized traits accounts for correlation.* Avoiding double counting of correlated traits

𝑊𝑍_𝑖_ *= matrix multiplication that scales each element of* 𝑍_𝑖_ *by its corresponding weight* 𝑊_𝑖_ .

𝑒_𝑖𝑗_*= residual (can be ignore)*

WHGCAMT is the overall strength of the GCA profile of an inbred, after scaling traits, applying weights for the important traits and corrected for correlation between traits. In the present study, non-negative weights reflecting breeding priorities (e.g., sum of 1) were picked. Under *Striga* infestation, weights of 0.3 for GYD, 0.15 each for SDS, ESP, EASP and EPP and 0.1 for ASI were applied. Under optimum environment, weights of 0.3 for GYD, 0.2 each for PASP and EASP and 0.1 each for HCV, ASI and EPP were used.

The 1696 SNP dosage format were used to generate Gowers’ matrix using the daisy function in the Gower package (47) implemented in R across *Striga*-infestation and optimum environments. The dissimilarity matrix was based on equation (6).

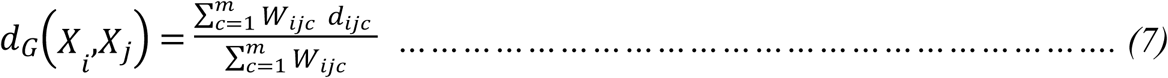

*Where: dijc is a dissimilarity measure between the i-th and j-th genotypes by the c-th variable (c = 1, …, m), and wijc takes the value zero, if either the i-th or the j-th genotype by the c-th variable is missing; otherwise, it takes the value one*.

Similarly, HSGCA and HGCAMT estimates were used to generate Euclidean distance matrices while WHGCAMT estimate were used to generate the inverse covariance matrix (Mahalanobis-like matrix). The matrices generated were subjected for hierarchical clustering using the average linkage method in the package dendextend (48) implemented in R. The optimal number clusters were identified through the Silhouette K-means analysis for a range of K from 2 to 10 in the package NbClust implemented in R (49). The final dendrogram was viewed using the package dendextend (48). The members within the same cluster were identified based on a color to distinguish them from members of another cluster. Analysis of molecular variance (AMOVA) based on the optimum number of clusters identified from the SNP was used to partition the total variation between and within clusters using the adegenet and poppr packages built in the R (50,51). The matrices derived from SNP markers and from the HSGCA, HGCAMT, and WHGCAMT estimates were converted from wide to long format to align with their corresponding half-diallel hybrids. Mid-parents, better-parents and economic heterosis were estimated following equations 9, 10 and 11, respectively. The Mean grain yield of the hybrids (GYD), SCA effects (SCA), mid-parents (MPH), better-parent (BPH), economic (ECH) heterosis, the converted matrices of the HSCA, HSGCA, HGCAMT, WHGCAMT and molecular marker (HMM) were subjected to correlation analysis using Performance Analytics in R. The GYD, SCA and ECH of the intra-group hybrids (all possible cross combinations resulting from the individuals of the same HG) and inter-group hybrids (all possible cross combinations made from the individuals of different HGs) of each heterotic grouping method were compared using a t-test. Similarly, the GYD, SCA, and ECH of the intra- and inter-group hybrids of all heterotic grouping methods were compared using ANOVA and a multiple t-test was used to compare each pair of HG. The breeding efficiency (BE) of each of the heterotic grouping method was calculated under *Striga*-infested and *Striga*-free environments using equation 13, modified version of equation 12 (52).

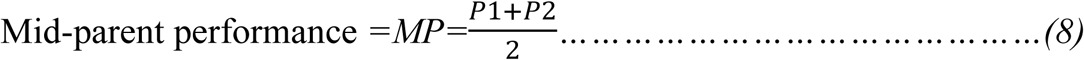

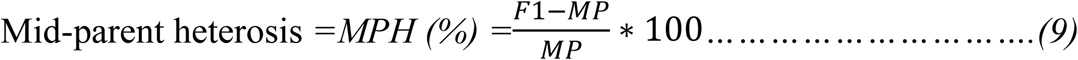

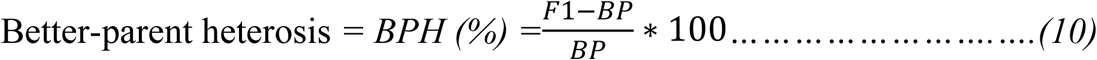

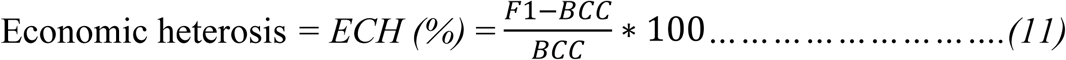

where MP= Mid-parent; P_1_= parent 1, P_2_=parent 2, BPH= better parent heterosis; BP=better parent; F_1_= Mean grain yield of the hybrid, ECH = economic heterosis; BCC= best commercial check

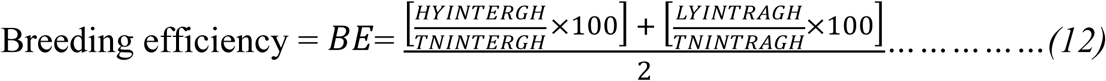

*where BE = breeding efficiency, HYINTERGH is the number of high-yielding inter-heterotic group hybrids, TNINTERGH is the total number of inter-heterotic group hybrids, LYINTRAGH is the number of low-yielding intra-heterotic group hybrids, and TNINTRAGH is the total number of intra-heterotic group hybrids*.

This procedure involved dividing the total number of hybrids into two major groups (intra- and inter-group hybrids) for each method. These two groups were subsequently divided into heterotic hybrids (hybrids that had yield greater than the best commercial check. i.e. high yielding hybrids), and non-heterotic hybrids (hybrids that had yield less than the lowest commercial check. i.e. low yielding hybrids). Therefore, the BE were estimated as follows:

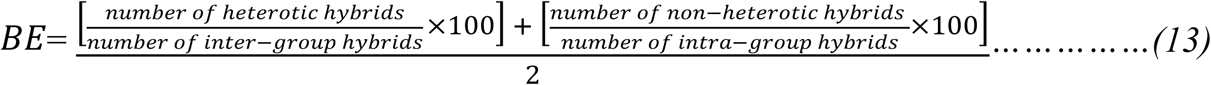

## Results

### Mean squares of the analysis of variance (ANOVA)

Under artificial *Striga* infestation, Environment (E) and genotype (G) mean squares were highly significant (p < 0.01) for all traits measured except for ASI. The GCA and SCA mean squares were also highly significant (p < 0.01) for all traits except the SCA mean squares for ASI. The environment × genotype (E × G) interaction was highly significant (p < 0.01) for GYD, SDS, EASP, PHT, and EHT, but not for ESP, EPP, and ASI. Additionally, the E × GCA mean square was also highly significant (p < 0.01) for all the measured traits. However, the E × SCA mean squares were not significant except for GYD, SDS, and EASP (Table 1). The combined ANOVA across Striga-free environments revealed highly significant (p < 0.01) mean squares for all sources of variation for all traits studied except for E × G and E × GCA mean squares for HCV. However, the E × SCA mean squares were not significant except for GYD, SDS, and EASP (Table 2).

**Table 1.** Mean squares from the combined ANOVA for grain yield and other *Striga*-adaptive traits of maize hybrids evaluated under artificial *Striga* infestation in Abuja and Mokwa

| Source | DF | GYD | SDS | ESP | EASP | PHT | EHT | EPP | ASI |
| --- | --- | --- | --- | --- | --- | --- | --- | --- | --- |
| Environment (E) | 1 | 356708372.38** | 31.36** | 1850.08** | 159.87** | 308936.85** | 66719.27** | 4.37** | 441.65** |
| Genotype (G) | 299 | 3545383.13** | 2.77** | 328.27** | 2.31** | 763.96** | 269.02** | 0.04** | 0.50ns |
| GCA | 24 | 28384410.48** | 21.72** | 1417.53** | 14.98** | 6994.61** | 2432.88** | 0.15** | 1.24** |
| SCA | 275 | 1377613.47** | 1.11** | 233.20** | 1.20** | 220.19** | 80.18** | 0.03** | 0.44ns |
| $E \times G$ | 299 | 708130.79** | 0.95** | 197.73ns | 0.67** | 107.66* | 41.58* | 0.02ns | 0.47ns |
| $E \times GCA$ | 24 | 2702231.69** | 3.90** | 626.62** | 1.68** | 245.63* | 83.92** | 0.04** | 0.92** |
| $E \times SCA$ | 275 | 534100.17** | 0.69** | 160.30ns | 0.58* | 95.62ns | 37.88ns | 0.02ns | 0.43ns |
| Residual | 534 | 440416.55 | 0.58 | 177.73 | 0.48 | 90.17 | 35.48 | 0.02 | 0.46 |

\*, significant at $p < 0.05$ ; \*\*, highly significant at $p < 0.01$ ; ns: not significant; DF: degrees of freedom; GYD: grain yield; SDS: *Striga* damage symptom; ESP: number of emerged *Striga* plants; PHT: plant height; EHT: ear height; EASP: ear aspect; EPP: ears per plant; ASI: anthesis silking interval; GCA: general combining ability; SCA: specific combining ability

**Table 2.** Mean squares from the combined ANOVA for grain yield and other agronomic traits of maize hybrids evaluated under optimum environments in Abuja and Mokwa.

| Source | DF | GYD | PASP | EASP | PHT | EHT | HCV | ASI | EPP |
| --- | --- | --- | --- | --- | --- | --- | --- | --- | --- |
| Environment (E) | 1 | 1277054741.69** | 115.94** | 266.96** | 53961.84** | 59840.56** | 186.44** | 128.71** | 0.13** |
| Genotype (G) | 299 | 7943923.12** | 2.68** | 2.55** | 943.52** | 420.98** | 0.93* | 0.33** | 0.04** |
| GCA | 24 | 76679958.52** | 21.89** | 18.23** | 8232.04** | 3841.29** | 1.67** | 0.81** | 0.20** |
| SCA | 275 | 1945141.85** | 1.00** | 1.18** | 307.43** | 122.48** | 0.86** | 0.29** | 0.03** |
| E × G | 299 | 1341583.42** | 0.73** | 0.87** | 234.01** | 88.70ns | 0.75ns | 0.24** | 0.02* |
| E × GCA | 24 | 3992795.05** | 1.32** | 1.29** | 341.23** | 134.50** | 1.02ns | 0.31* | 0.03** |
| E × SCA | 275 | 1110204.96** | 0.68** | 0.83** | 224.66** | 84.71ns | 0.73ns | 0.23** | 0.02* |
| Residual | 534 | 675176.68 | 0.50 | 0.64 | 168.65 | 78.31 | 0.69 | 0.18 | 0.02 |

\*, significant at $p < 0.05$ ; \*\*, highly significant at $p < 0.01$ ; ns: not significant; df: degrees of freedom; GYD: grain yield; PHT: plant height; EHT: ear height; PASP: plant aspect; EASP: ear aspect; HCV: husk cover; EPP: ears per plant; ASI: anthesis silking interval; GCA: general combining ability; SCA: specific combining ability

### Genetic diversity indices

The observed heterozygosity (Ho) ranged from 0.00 to 0.44, with an average of 0.04 (Table 3). However, the majority of loci attained homozygosity, with nearly 95% exhibiting Ho < 0.01 (Figure 1A). The expected heterozygosity (He) varied between 0.09 and 0.50, with an average of 0.37 (Table 3). Minor allele frequency (MAF) ranged from 0.05 to 0.50, with a mean of 0.28 and evenly distributed across the markers (Figure 1B). Polymorphic information content (PIC) of the marker varied from 0.01 to 0.50, with an average of 0.37 (Table 3). However, majority of the makers have PIC >0.4 (Figure 1C). The distribution of the 1,696 high-quality SNPs revealed that 161, 159, 179, 177, 216, 148, 154, 171, 138 and 113 SNP markers were located on chromosomes 1 to 10, respectively (Figure 1D).

**Figure 1.**
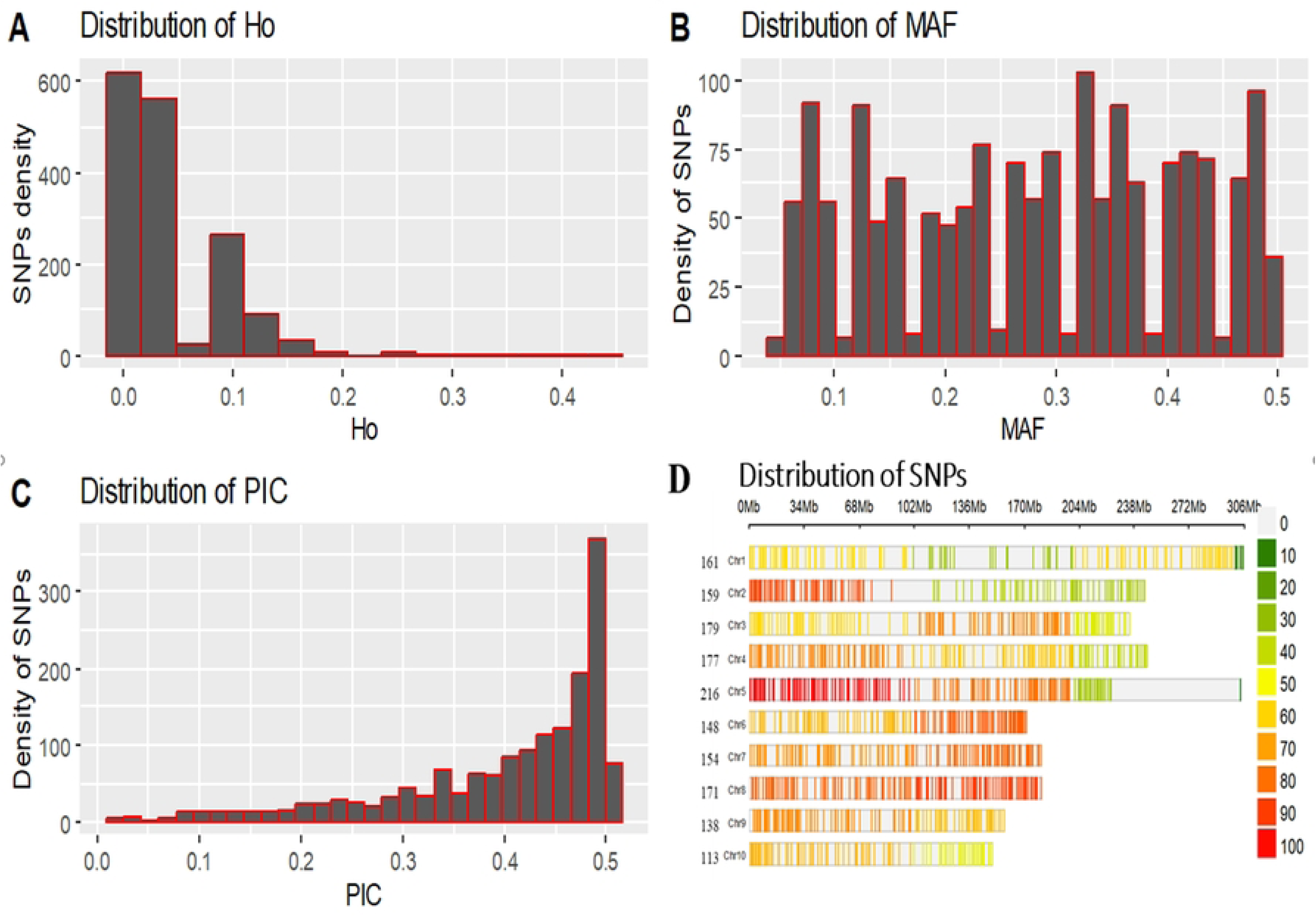
Bar plot showing the diversity indices of the SNP markers used in the study including (A) observed heterozygosity; (B) minor alleles frequency; (C) polymorphic information content of the 1696 SNP markers and (D) distribution of the SNPs. *Ho: observed heterozygosity; MAF: minor allele frequency; PIC: polymorphic information content*

**Table 3.** Genetic diversity indices of the 25 inbred lines based on 1696 SNP markers

| Statistic | $H_o$ | $H_e$ | MAF | PIC |
| --- | --- | --- | --- | --- |
| Minimum | 0.00 | 0.09 | 0.05 | 0.10 |
| Maximum | 0.44 | 0.50 | 0.50 | 0.50 |
| Average | 0.04 | 0.37 | 0.28 | 0.40 |

$H_o$ = observed heterozygosity, $H_e$ = expected heterozygosity, MAF = minor allele frequency, PIC = polymorphic information content.

### Analysis of molecular variance (AMOVA)

The AMOVA results showed significant genetic differences (Φ = 0.38, *P = 0.034*) between clusters. Variation between clusters accounts for 38% of the total genetic variance. However, 62 % of the genetic variation was observed among the individuals within the clusters (Table 4).

**Table 4.** Analysis of molecular variance (AMOVA) among and within 3 clusters (K=3) of 25 inbred lines based on 1696 SNP markers

| Source of variation | df | Sum Squares | Mean Squares | Estimated variance | % variance | Phi ($\Phi$) | P-value |
| --- | --- | --- | --- | --- | --- | --- | --- |
| Between clusters | 2 | 6073.11 | 3036.55 | 325.41 | 37.72 | 0.38 | 0.034 |
| Within cluster | 22 | 11822.97 | 537.41 | 537.41 | 62.29 | 0.62 |  |
| Total | 24 | 17896.08 | 745.67 | 862.82 | 100.00 |  |  |

DF: degrees of freedom

### Mode of gene action

The relative importance of GCA and SCA effects was assessed by expressing the ratio of GCA effects to the total genetic effects. The closer the ratio to unity, the greater the predictability based on GCA alone (45). GCA effects accounted for 95 % and 97 % of the total genetic effects for GYD under Striga infestation and optimum environments, respectively, (Figure 2). Generally, the GCA effects modulated the inheritance of all traits under *Striga* infestation and in optimal environments.

**Figure 2.**
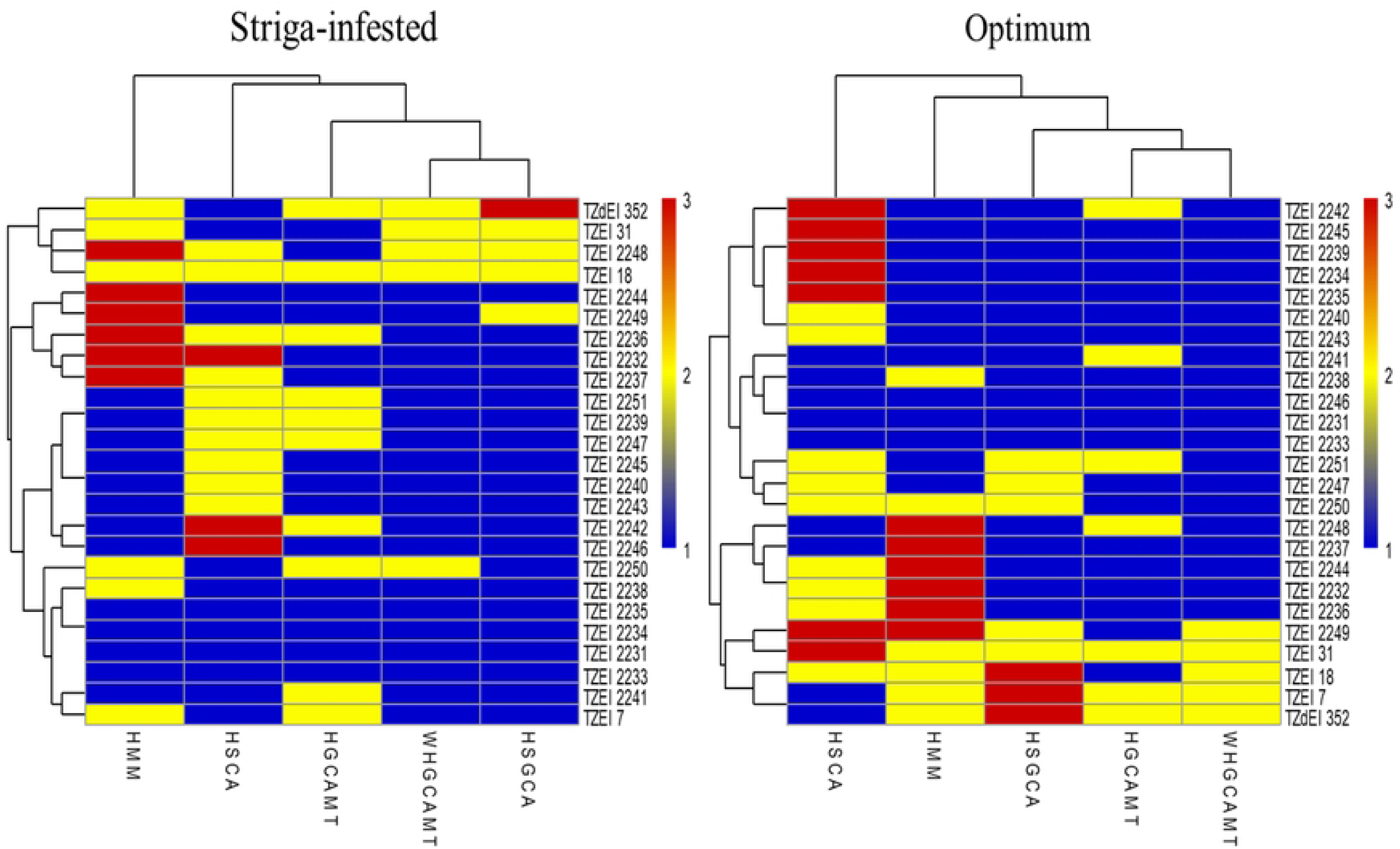
Proportionate contribution of GCA and SCA variances to the total genetic effects (twice the GCA plus the SCA effects) of GYD and other *Striga*-adaptive traits of 25 inbred lines evaluated under *Striga* infestation and optimum environments. *GYD: grain yield; SDS: Striga damage symptom; ESP: number of emerged Striga plants; PHT: plant height; EHT: ear height; EASP: ear aspect; EPP: ears per plant; ASI: anthesis silking interval; PASP: plant aspect; HCV: husk cover; GCA: general combining ability; SCA: specific combining ability*

### General combining ability effects of the parental lines

Under *Striga* infestation, significant (p ≤ 0.001) positive GCA effects for GYD were observed in the inbreds TZEI 2248, TZEI 2249, TZEI 2250, TZEI 2251, TZEI 18 and TZdEI 352.

In contrast, significant (p ≤ 0.001) negative GCA effects for GYD were detected in the inbreds TZEI 2231, TZEI 2232, TZEI 2233, TZEI 2234, TZEI 2235, TZEI 2239, TZEI 2240, TZEI 2241, TZEI 2242, TZEI 2243, TZEI 2245, TZEI 2247, TZEI 7 and TZEI 31 (Table 5). The majority of the inbred lines with significant (p ≤ 0.001) positive GCA effects for GYD possessed significant positive GCA effects for EPP and significant negative GCA effects for SDS, ESP and EASP (Table 5). Similarly, under optimum environment (Table 6), inbred lines TZEI 2247, TZEI 2248, TZEI 2249, TZEI 2250, TZEI 2251, TZEI 7, TZEI 18, TZEI 31 and TZdEI 352 had significant (p ≤ 0.001) positive GCA effects for GYD, while TZEI 2231, TZEI 2232, TZEI 2233, TZEI 2234, TZEI 2235, TZEI 2236, TZEI 2237, TZEI 2238, TZEI 2239, TZEI 2240, TZEI 2242, TZEI 2243, TZEI 2245 and TZEI 2246 recorded significant (p ≤ 0.001) negative GCA effects for GYD. Similar to the result observed under *Striga* infestation, the majority of the inbred lines exhibiting significant positive GCA effects for GYD also demonstrated significant (p≤0.001) positive GCA effects for EPP and significant (p≤0.001) negative GCA effects for PASP and EASP (Table 6).

**Table 5.** GCA effects of inbred parents for grain yield and other *Striga*-adaptive traits evaluated in Abuja and Mokwa under *Striga*-infested environments in Nigeria

| Entry | Line | GYD | SDS | ESP | EASP | PHT | EHT | EPP | ASI |
| --- | --- | --- | --- | --- | --- | --- | --- | --- | --- |
| 1 | TZEI 2231 | -391.84** | 0.41** | -1.51ns | 0.24** | -5.06** | -2.89** | 0.00ns | 0.02ns |
| 2 | TZEI 2232 | -484.30** | 0.18** | -2.89** | 0.37** | -5.22** | -2.32** | 0.01ns | 0.05ns |
| 3 | TZEI 2233 | -213.08** | 0.18** | 0.78ns | 0.10ns | -8.92** | -5.14** | 0.03** | 0.04ns |
| 4 | TZEI 2234 | -220.00** | 0.31** | 1.29ns | 0.08ns | -10.50** | -6.26** | -0.01ns | 0.11* |
| 5 | TZEI 2235 | -490.47** | 0.57** | 3.00** | 0.37** | -11.76** | -6.03** | -0.01ns | 0.17** |
| 6 | TZEI 2236 | 74.13ns | -0.20** | -1.86ns | -0.05ns | -1.19ns | -1.21** | 0.03* | -0.11* |
| 7 | TZEI 2237 | -79.06ns | -0.04** | 0.93ns | 0.05ns | -1.78* | -1.13** | 0.03** | -0.04ns |
| 8 | TZEI 2238 | -354.72** | 0.39** | 3.29** | 0.28** | -6.06** | -4.11** | 0.00ns | -0.04ns |
| 9 | TZEI 2239 | -26.95ns | -0.12** | -1.68ns | 0.01ns | -0.03ns | -0.19ns | 0.03** | -0.12* |
| 10 | TZEI 2240 | -292.97** | 0.05 | 3.64** | 0.07ns | -3.20** | -2.56** | 0.02* | 0.08ns |
| 11 | TZEI 2241 | -651.40** | 0.64** | -5.05** | 0.49** | -4.61** | -2.33** | -0.08** | -0.07ns |
| 12 | TZEI 2242 | -645.48** | 0.52** | -5.87** | 0.49** | -5.36** | -4.57** | -0.05** | 0.06ns |
| 13 | TZEI 2243 | -448.21** | 0.38** | -2.39* | 0.33** | -13.02** | -6.90** | 0.01ns | 0.01ns |
| 14 | TZEI 2244 | 32.29ns | 0.11ns | 0.09ns | 0.14** | -9.81** | -3.43** | -0.02* | 0.16** |
| 15 | TZEI 2245 | -143.27** | -0.07ns | 4.66** | 0.14** | -3.10** | -1.76** | -0.01ns | 0.04ns |
| 16 | TZEI 2246 | 46.77ns | -0.20** | 4.79** | -0.08ns | 3.69** | -0.57ns | -0.01ns | 0.20** |
| 17 | TZEI 2247 | -163.99** | 0.18** | -6.74** | 0.07ns | 10.31** | 9.41** | -0.05** | 0.16** |
| 18 | TZEI 2248 | 725.33** | -0.63** | -5.32** | -0.42** | 9.29** | 2.55** | 0.05** | 0.06ns |
| 19 | TZEI 2249 | 511.82** | -0.49** | 5.62** | -0.35ns | 4.87** | 2.05** | 0.03** | -0.16** |
| 20 | TZEI 2250 | 1063.16** | -0.93** | -1.37ns | -0.76** | 15.54** | 8.63** | 0.08** | -0.25** |
| 21 | TZEI 2251 | 513.05** | -0.56** | -1.30ns | -0.56** | 6.25** | 3.37** | 0.04** | -0.13** |
| 22 | TZEI 7 | -32.54* | 0.23** | -1.39ns | 0.23** | 3.86** | 3.09** | -0.10** | 0.02ns |
| 23 | TZEI 18 | 824.19** | -0.40** | -1.2ns4 | -0.43** | 11.00** | 6.23** | -0.02* | -0.06ns |
| 24 | TZEI 31 | -611.71** | 0.67** | 9.67** | 0.33** | 4.88** | 3.49** | -0.02* | -0.05ns |
| 25 | TZdEI 352 | 14 59.25** | -1.19** | 0.84ns | -1.14** | 19.92** | 12.57** | 0.04** | -0.16** |
|  | SE | 47.94 | 0.06 | 0.96 | 0.050 | 0.67 | 0.43 | 0.01 | 0.05 |

\*: significant at $p < 0.05$ ; \*\*: highly significant at $p < 0.01$ ; ns: not significant; GYD: grain yield; SDS: *Striga* damage symptom; ESP: number of emerged *Striga* plants; PHT: plant height; EHT: ear height; EASP: ear aspect; EPP: ears per plant; ASI: anthesis silking interval; SE: standard error

**Table 6.** GCA effects of inbred parents for grain yield and other agronomic traits evaluated in Abuja and Mokwa under optimum environments in Nigeria

| Entry | Line | GYD | PASP | EASP | PHT | EHT | HCV | ASI | EPP |
| --- | --- | --- | --- | --- | --- | --- | --- | --- | --- |
| 1 | TZEI 2231 | -809.22** | 0.50** | 0.30** | -8.33** | -5.09** | 0.01ns | 0.05ns | -0.06** |
| 2 | TZEI 2232 | -938.29** | 0.44** | 0.54** | -4.59** | -3.77** | 0.01ns | 0.06* | -0.05** |
| 3 | TZEI 2233 | -836.43** | 0.29** | 0.37** | -9.67** | -5.92** | 0.15ns | 0.06* | -0.03** |
| 4 | TZEI 2234 | -824.30** | 0.50** | 0.46** | -11.63** | -8.88** | 0.05ns | 0.18** | -0.04** |
| 5 | TZEI 2235 | -709.81** | 0.52** | 0.36** | -10.08** | -7.02** | 0.05ns | 0.08** | -0.06** |
| 6 | TZEI 2236 | -532.88** | 0.26** | 0.24** | -2.94** | -2.96** | 0.13* | -0.01ns | -0.02** |
| 7 | TZEI 2237 | -476.77** | 0.17** | 0.41** | -2.36* | -2.90** | -0.02ns | 0.01ns | -0.05** |
| 8 | TZEI 2238 | -537.62** | 0.36** | 0.36** | -4.12** | -3.23** | 0.22** | 0.03ns | -0.04** |
| 9 | TZEI 2239 | -679.38** | 0.32** | 0.29** | -5.85** | -4.42** | 0.08ns | 0.11** | -0.02** |
| 10 | TZEI 2240 | -558.38** | 0.24** | 0.30** | -3.23** | -4.14** | -0.07ns | 0.03ns | -0.02* |
| 11 | TZEI 2241 | -51.77ns | -0.01ns | -0.03ns | -2.18** | -1.55* | -0.28** | -0.14** | 0.00ns |
| 12 | TZEI 2242 | -243.35** | 0.03ns | 0.02ns | -3.75** | -2.59** | -0.13* | -0.06ns | -0.03** |
| 13 | TZEI 2243 | -970.60** | 0.64** | 0.53** | -16.04** | -8.98** | 0.18** | 0.05ns | -0.06** |
| 14 | TZEI 2244 | -9.85ns | 0.16** | -0.05ns | -11.71** | -3.86** | 0.14* | -0.01ns | 0.04** |
| 15 | TZEI 2245 | -553.27** | 0.21** | 0.26** | -2.96** | 0.02ns | 0.18** | -0.09** | -0.02* |
| 16 | TZEI 2246 | -188.18** | 0.06ns | 0.05ns | 2.21* | -1.91** | -0.02ns | -0.01ns | 0.00ns |
| 17 | TZEI 2247 | 622.99** | -0.35** | -0.31** | 12.04** | 13.01** | -0.02ns | 0.13** | 0.03** |
| 18 | TZEI 2248 | 125.53* | -0.23** | -0.25** | 9.48** | 3.17** | -0.14* | -0.01ns | 0.02ns |
| 19 | TZEI 2249 | 702.74** | -0.18** | -0.43** | 3.40** | 3.11** | 0.00ns | 0.00ns | 0.05** |
| 20 | TZEI 2250 | 496.62** | -0.22** | -0.30** | 7.21** | 5.75** | 0.03ns | -0.12** | 0.07** |
| 21 | TZEI 2251 | 210.31** | -0.34** | -0.25** | 7.77** | 3.21** | -0.01ns | -0.11** | 0.02* |
| 22 | TZEI 7 | 1971.59** | -0.84** | -0.64** | 14.59** | 8.60** | -0.26** | -0.11** | 0.06** |
| 23 | TZEI 18 | 1618.31** | -0.66** | -0.58** | 11.93** | 9.62** | 0.07ns | -0.13** | 0.06** |
| 24 | TZEI 31 | 900.29** | -0.41** | -0.40** | 11.23** | 7.83** | -0.19** | -0.12** | 0.07** |
| 25 | TZdEI 352 | 2271.76** | -1.46** | -1.26** | 19.58** | 12.88** | -0.15* | 0.15** | 0.08** |
|  | SE | 393.35 | 0.05 | 0.06 | 0.94 | 0.64 | 0.06 | 0.03 | 0.01 |

\*: significant at $p < 0.05$ ; \*\*: highly significant at $p < 0.01$ ; ns: not significant; GYD: grain yield; PHT: plant height; EHT: ear height; PASP: plant aspect; EASP: ear aspect; HCV: husk cover; EPP: ears per plant; ASI: anthesis silking interval; SE: standard error

### Specific combining ability effects of the crosses

The SCA effects of the crosses evaluated under *Striga* infestation for grain yield and other *Striga*-adaptive traits are presented in supplementary Table S3. The highest significant positive SCA effects for GYD were observed in the crosses between TZEI 2238 × TZEI 2240 (2036.03, *p ≤ 0.001*) followed by TZEI 2247 × TZdEI 352 (1579.90, *p ≤ 0.001*), TZEI 2242 × TZEI 18 (1519.98, *p ≤ 0.001*), TZEI 2237 × TZEI 2244 (1367.76, *p ≤ 0.001*), TZEI 2241 × TZEI 7 (1188.72, *p≤ 0.001*), TZEI 2244 × TZdEI 352 (1187.89, *p ≤ 0.001*) and TZEI 2232 × TZEI 2249 (1186.98, *p ≤ 0.001*).

The lowest significant negative SCA effects for GYD were observed in the crosses between TZEI 2245 × TZEI 2246 (−2060.95, *p≤0.001*), followed by TZEI 2249 × TZEI 7 (−2007.96, *p≤0.001*), TZEI 2244 × TZEI 2251 (−1450.23, *p≤0.001*), TZEI 31 × TZdEI 352 (−1427.38, *p≤0.001*), TZEI 2231 × TZEI 2247 (−1219.47, *p≤0.001*) and TZEI 2239 × TZEI 2249 (−1216.07, *p≤0.001*). In summary, of the 300 crosses evaluated under *Striga* infestation, approximately 45 and 40 crosses displayed significant positive and negative SCA effects for GYD, respectively. Similar to the observed in inbred lines with significant positive GCA effects for grain yield, majority of the crosses showed significant positive SCA effects for GYD, recorded significant positive SCA effects for EPP and significant negative SCA effects for SDS, ESP and EASP (Table S2). The estimated SCA effects under optimum environment showed that approximately 30 and 40 crosses displayed significant (*p≤0.001*) positive and negative SCA effects for GYD, respectively, (Table S3). The highest significant (*p≤0.001*) positive SCA effects were recorded in the crosses between TZEI 2238 × TZEI 2240 (2755.06, *p≤0.001*), followed by TZEI 2237 × TZEI 2247 (2020.99, *p≤0.001*), TZEI 2240 × TZEI 7 (1646.55, *p≤0.001*) and TZEI 2235 × TZdEI 352 (1596.32, *p≤0.001*). The lowest significant negative SCA effects were detected in the crosses between TZEI 2241 × TZEI 2242 (−1875.26, *p≤0.001*), followed by TZEI 7 × TZEI 31 (−1699.78, *p≤0.001*), TZEI 7 × TZEI 31 (−1699.78, *p≤0.001*), TZEI 18 × TZdEI 352 (−1674.35, *p≤0.001*) and TZEI 2237 × TZEI 2241 (−1572.57, *p≤0.001*). Seven of the 30 hybrids showed significant positive SCA effects for grain yield under *Striga* infestation environment

### Grain yield performance of the parental lines, their crosses and checks evaluated across *Striga* infestation and optimum environments

The mean GYD of the parental lines evaluated under *Striga* infestation ranged from 166.0 kg/ha for TZEI2234 to 1760 kg/ha for TZEI18 with an average of 1017.9 kg/ha (Table S4). Similarly, the mean grain yield performance of the hybrids evaluated under *Striga* infestation varied from 838.9 kg/ha for TZEI2245 × TZEI2246 to 5770.8 kg/ha for TZEI2248 × TZEI18 with an average of 2860.4 kg/ha (Table S5). Under optimum environment, the mean grain yield of the hybrid varied from 1491.8 kg/ha for TZEI2233 × TZEI2234 to 7242.6 kg/ha for TZEI2241 × TZdEI352 with an average of 4049.1 kg/ha (Table S6).

### Heterotic grouping of the parental lines

Both under artificial *Striga* infestation and optimum environments, Silhouette K-means analysis for a range of K set from two to ten indicated that the optimum clusters were two (i.e., k=2) based on the HGCAMT and WHGCAMT methods (Figure S1a and S1b). However, the optimal number of clusters was three based on the molecular marker (HMM) and HSGCA methods (Figure S1c and S1d). Similarly, the inbred lines were classified into three HGs based on HSCA.

Under *Striga* Striga-infested environment, the results of the dendrogram constructed based on the HGCAMT placed 10 (TZEI 2236, TZEI 2239, TZEI 2241, TZEI 2242, TZEI 2247, TZEI 2250, TZEI 2251, TZEI 7, TZEI 18 and TZdEI 352) and 15 inbred lines into HGs 1 and 2, respectively (Figure S1a). The WHGCAMT method placed 20 inbred lines into HG 1 and 5 other inbred lines including TZEI 2248, TZEI 2250, TZEI 18, TZEI 31 and TZdEI 352 in another HG (Figure S1b). Using the molecular marker approach, the inbred lines were classified into HGs 1, 2, and 3 with corresponding numbers of 13, 6 and 6 (Figure S1c). The HSGCA approach classified 20 inbred lines into HG 1 and 4 inbred lines (TZEI 2248, TZEI 2249, TZEI 18 and TZEI 31) into HG 2 and placed TZdEI 352 into a different HG (Figure S1d). Similarly, the HSCA placed 12, 10 and 3 inbred lines into HGs 1, 2, and 3, respectively (Table S7).

Under optimum environment, HGCAMT approach placed 18 and 7 (TZEI 2241, TZEI 2242, TZEI 2248, TZEI 2251, TZEI 7, TZEI 31 and TZdEI 352) inbred lines into HGs 1 and 2, respectively, and WHGCAMT method placed 20 inbred lines into HG 1 and 5 other inbred lines (TZEI 2231, TZEI 2249, TZEI 7, TZEI 18, TZEI 31 and TZdEI 352) in the second HG. The HSGCA approach placed 17 inbred lines into HG 1 and 5 inbred lines comprising TZEI 2247, TZEI 2249, TZEI 2250, TZEI 2251 and TZEI 31 into HG 2 and placed TZEI 7, TZEI 18 and TZdEI 352 into the third HG. Similarly, 9 inbred lines each were placed in HGs 1 and 2 by HSCA and 7 inbred lines into HG 3 (Table S7).

### Comparison of effectiveness of the heterotic grouping methods

When inbred lines were placed into HGs, the five classification methods showed similar trends but were not identical to one another. The detailed description of the placement of the 25 inbred lines into different HGs and the association between clusters under *Striga* infestation and optimum environments was presented in Figure 3. In summary, the number of clusters observed for each method under artificial *Striga* infestation and optimum environments are the same. Each HGCAMT and WHGCAMT placed the inbred lines into 2 HGs while HSCA, HSGCA and HMM placed them into 3 HGs. The highest consistency of the placement of the inbred lines into a similar group under *Striga* infestation and optimum environments were observed in WHGCAMT followed by HSGCA, HGCAMT and HSCA (Figure 3). Of the 25 inbred lines, 21, 19, 18 and 15 were placed in the same group under *Striga* infestation and optimum environments by WHGCAMT, HSGCA, HGCAMT and HSCA, respectively (Figure 3). Additionally, comparable patterns were observed in the grouping of the 25 inbred lines by the different methods. However, the WHGCAMT shows close correspondence with HSGCA under both Striga-infested and optimal environments. Although WHGCAMT detected two HGs, HSGCA identified three; of the 25 parental lines, WHGCAMT and HSGCA agreed in the placement of 22 and 17 inbred parents into similar HGs under *Striga* infestation and optimum environments, respectively (Figure 3).

**Figure 3.**
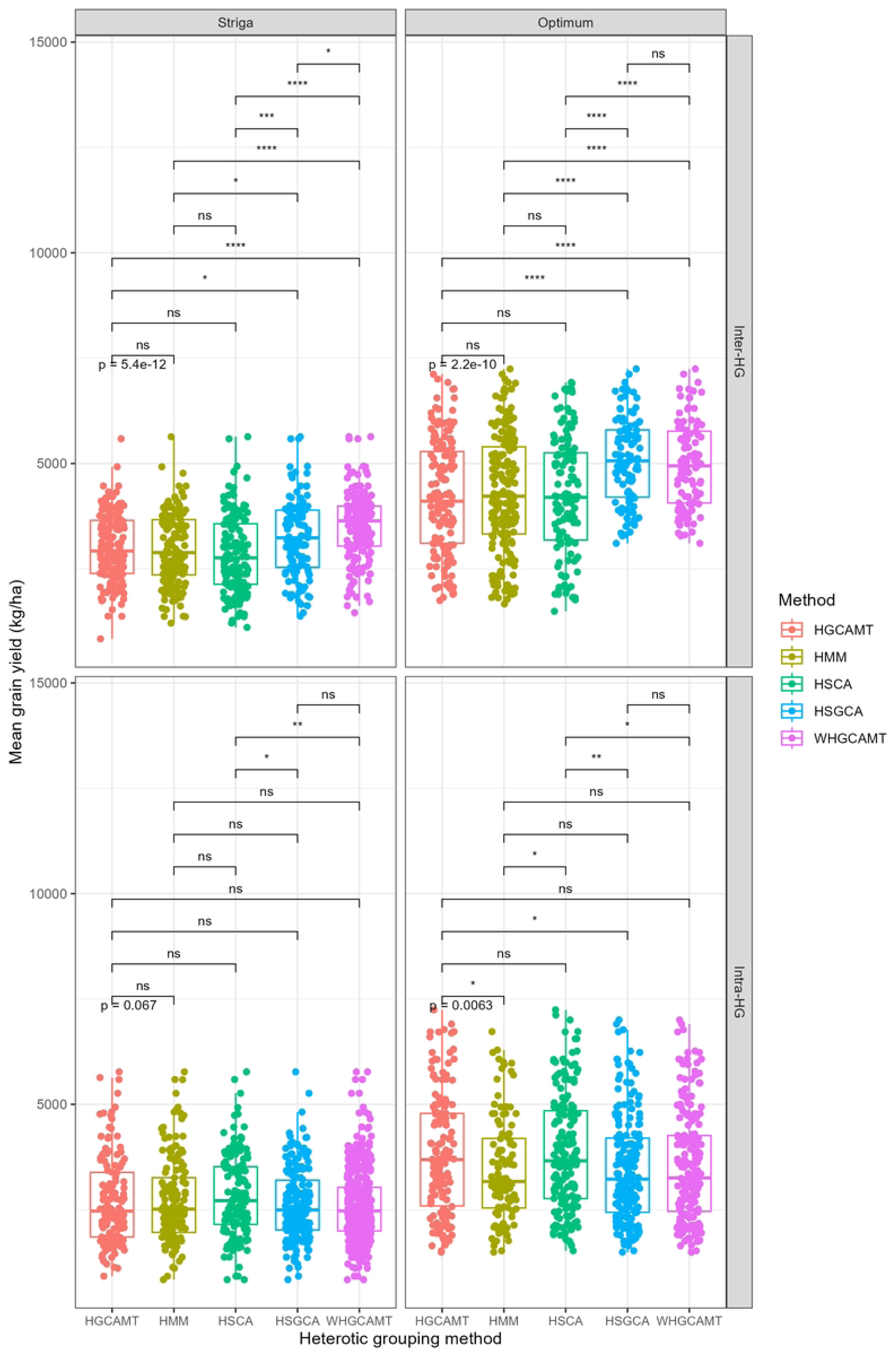
Pheatmap showing the relationship within and between the inbred lines and heterotic grouping methods. *HSCA: heterotic grouping based on SCA of grain yield; HSGCA: heterotic grouping based on SCA and GCA of grain yield; HMM: heterotic grouping based on molecular marker; HGCAMT: heterotic grouping based on GCA of multiple traits; WHGCAMT: weighted heterotic grouping based on GCA of multiple traits; <u>blue: HG1; cream: HG2; red: HG3</u>*

Under Striga infestation, the heterotic grouping methods HSCA, HSGCA, HMM, HGCAMT, and WHGCAMT had average grain yields (GYD) for intra- and inter-group crosses of 2857.2 and 2858.3, 2622.0 and 3242.5, 2735.7 and 2973.5, 2703.9 and 2999.8, and 2513.6 and 3333.1 kg/ha, respectively (Table 7).

**Table 7.** Average GYD, SCA and ECH of the intra and inter-group crosses produced by HSCA, HSGCA, HMM, HGCAMT and WHGCAMT heterotic grouping methods under *Striga*-infested and optimum environments

| Heterotic group | Measure | Hybrid type | Striga-infested |  |  | Optimum |  |  |
| --- | --- | --- | --- | --- | --- | --- | --- | --- |
|  |  |  | Maximum | Average | Minimum | Maximum | Average | Minimum |
| HSCA | GYD | Intra | 5770.8 | 2857.2 | 838.9 | 7242.6 | 4002.1 | 1491.8 |
|  |  | Inter | 5634.2 | 2858.3 | 1109.3 | 7242.6 | 4014.1 | 1491.8 |
|  | SCA | Intra | 1188.7 | -6 | -2060.9 | 2755.1 | 4.7 | -1875.3 |
|  |  | Inter | 2036 | 5.6 | -2008 | 2755.1 | 0.3 | -1875.3 |
|  | ECH | Intra | 250.3 | 72.9 | -49.1 | 20.5 | -33.4 | -75.2 |
|  |  | Inter | 242 | 74 | -32.7 | 20.5 | -33.2 | -75.2 |
| HSGCA | GYD | Intra | 5770.8 | 2622 | 838.9 | 7242.6 | 3994.6 | 1491.8 |
|  |  | Inter | 5634.2 | 3242.5 | 1374.6 | 7242.6 | 4095.6 | 1491.8 |
|  | SCA | Intra | 1367.8 | -34.2 | -2060.9 | 2755.1 | 10.3 | -1875.3 |
|  |  | Inter | 2036 | 55.8 | -2008 | 2755.1 | 12.3 | -1875.3 |
|  | ECH | Intra | 250.3 | 59.9 | -49.1 | 20.5 | -33.5 | -75.2 |
|  |  | Inter | 242 | 95.6 | -16.6 | 20.5 | -31.9 | -75.2 |
| HMM | GYD | Intra | 5770.8 | 2735.7 | 838.9 | 7242.6 | 3972.2 | 1491.8 |
|  |  | Inter | 5634.2 | 2973.5 | 1211.3 | 7242.6 | 4014.1 | 1491.8 |
|  | SCA | Intra | 2036 | -49.5 | -2060.9 | 2755.1 | 0.3 | -1875.3 |
|  |  | Inter | 1579.9 | 47 | -1450.2 | 2755.1 | 31.3 | -1875.3 |
|  | ECH | Intra | 250.3 | 66.3 | -49.1 | 20.5 | -33.2 | -75.2 |
|  |  | Inter | 242 | 80.2 | -26.5 | 20.5 | -33.9 | -75.2 |
| HGCAMT | GYD | Intra | 5770.8 | 2703.9 | 921.9 | 7242.6 | 4002.1 | 1491.8 |
|  |  | Inter | 5585.3 | 2999.8 | 838.9 | 7242.6 | 4034.9 | 1491.8 |
|  | SCA | Intra | 2036 | -129.1 | -2008 | 2755.1 | 4.7 | -1875.3 |
|  |  | Inter | 1520 | 119.2 | -2060.9 | 2755.1 | 3.6 | -1875.3 |
|  | ECH | Intra | 250.3 | 64.4 | -44 | 20.5 | -33.4 | -75.2 |
|  |  | Inter | 239 | 81.8 | -49.1 | 20.5 | -32.9 | -75.2 |
| WHGCAMT | GYD | Intra | 5770.8 | 2513.6 | 838.9 | 7242.6 | 4008.0 | 1491.8 |
|  |  | Inter | 5634.2 | 3333.1 | 1455.8 | 7242.6 | 4079.0 | 1491.8 |
|  | SCA | Intra | 1367.8 | -19.5 | -2060.9 | 2755.1 | 0.0 | -1875.3 |
|  |  | Inter | 2036 | 26.9 | -2008 | 2755.1 | 28.5 | -1875.3 |
|  | ECH | Intra | 250.3 | 53.4 | -49.1 | 20.5 | -33.3 | -75.2 |
|  |  | Inter | 242 | 101.3 | -11.6 | 20.5 | -32.1 | -75.2 |

HSCA: heterotic grouping based on SCA of grain yield; HSGCA: heterotic grouping based on SCA and GCA of grain yield; HMM: heterotic grouping based on molecular marker; HGCAMT: heterotic grouping based on GCA of multiple traits; WHGCAMT: weighted heterotic grouping based on GCA of multiple traits; GYD: grain yield; SCA: specific combining ability; ECH: economic heterosis

The average specific combining ability (SCA) effects for intra- and inter-group hybrids were −6 and 5.6 for HSCA, −34.2 and 55.8 for HSGCA, −49.5 and 47 for HMM, −129.1 and 119.2 for HGCAMT, and - 19.5 and 26.9 for WHGCAMT (Table 7). Corresponding average economic heterosis values for intra- and inter-group hybrids were 72.9 and 74 for HSCA, 59.9 and 95.6 for HSGCA, 66.3 and 80.2 for HMM, 64.4 and 81.8 for HGCAMT, and 53.4 and 101.3 for WHGCAMT, respectively (Table 7).

Under optimum environmental conditions, the average GYD for intra- and inter-group hybrids were 4002.1 and 4014.1 for HSCA, 3994.6 and 4095.6 for HSGCA, 3972.2 and 4014.1 for HMM, 4002.1 and 4034.9 for HGCAMT, and 4008.0 and 4079.0 kg/ha for WHGCAMT, respectively. The average SCA values for intra- and inter-group hybrids under optimum conditions were 4.7 and 0.3 for HSCA, 10.3 and 12.3 for HSGCA, 0.3 and 31.3 for HMM, 4.7 and 3.6 for HGCAMT, and 0.0 and 28.5 for WHGCAMT, respectively. The corresponding economic heterosis values were −33.4 and −33.2 for HSCA, −33.5 and −31.9 for HSGCA, −33.2 and −33.9 for HMM, −33.4 and −32.9 for HGCAMT, and −33.3 and −32.1 for WHGCAMT (Table 7).

Significant differences (*p < 0.05*) were detected in the grain yields of intra- and inter-group crosses produced by each heterotic grouping method under both *Striga* infestation and optimum environments, except for the HSCA method under Striga infestation (Figure 4). Similar trends were observed for economic heterosis (Figure S2). Conversely, SCA values under *Striga* infestation were not significantly different between intra- and inter-group crosses for any method, except HGCAMT. However, under optimum conditions, SCA values were significant (p < 0.05) for all methods except HGCAMT and HSCA (Figure S3).

**Figure 4.**
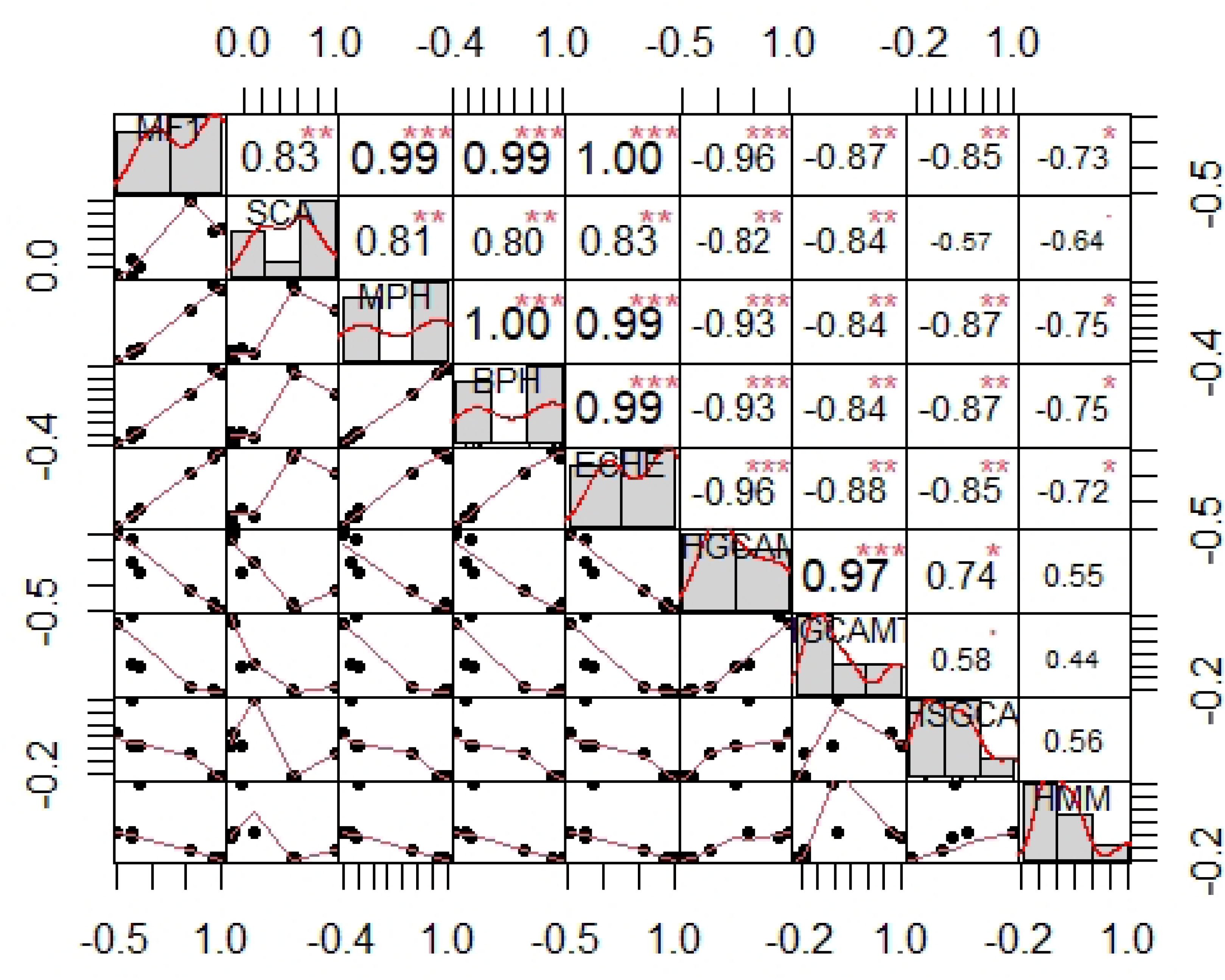
Box plot showing pair comparison between intra- and inter-group crosses for GYD of five different heterotic grouping methods under Striga-infested and optimum environments. *HSCA: heterotic grouping based on SCA of grain yield; HSGCA: heterotic grouping based on SCA and GCA of grain yield; HMM: heterotic grouping based on molecular marker; HGCAMT: heterotic grouping based on GCA of multiple traits; WHGCAMT: weighted heterotic grouping based on GCA of multiple traits*

Under *Striga* infestation, pair-wise t-test indicated significant (*p<0.05*) differences for GYD of the inter-group hybrids for HGCAMT and HSGCA, HGCAMT and WHGCAMT, HMM and HSGCA, HMM and WHGCAMT, HSCA and HSGCA, HSCA and WHGCAMT and HSGCA and WHGCAMT (Figure 5).

**Figure 5.**
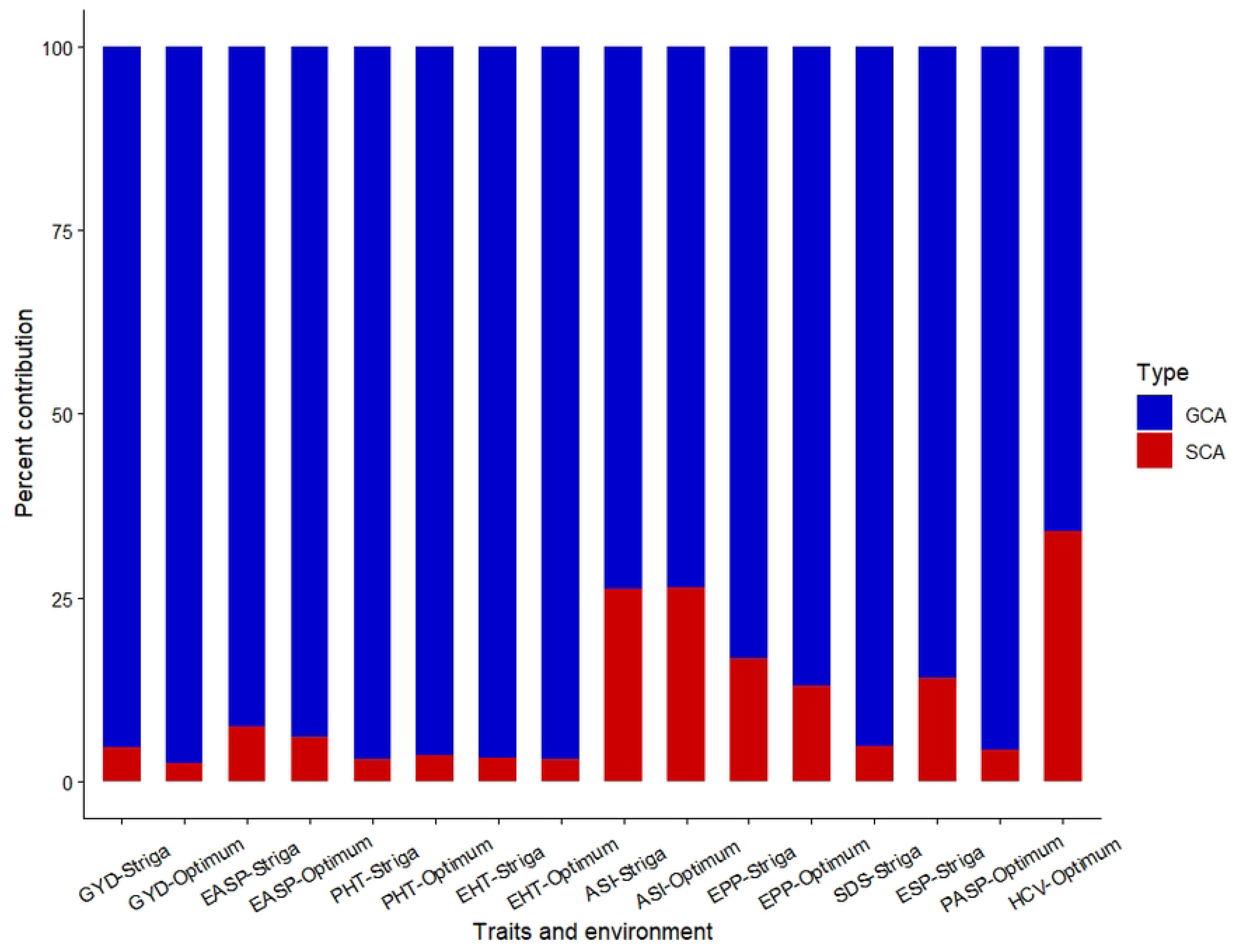
Box plot showing pair comparison among the heterotic grouping methods for GYD produced by intra- and inter-group crosses under *Striga*-infested and optimum environments. *HSCA: heterotic grouping based on SCA of grain yield; HSGCA: heterotic grouping based on SCA and GCA of grain yield; HMM: heterotic grouping based on molecular marker; HGCAMT: heterotic grouping based on GCA of multiple traits; WHGCAMT: weighted heterotic grouping based on GCA of multiple traits*

The GYD of the intra-group hybrids of HSCA and HSGCA and HSCA and WHGCAMT were also significant (p<0.05). Under optimum environment, significant differences were detected between the GYD produced in inter-group hybrids of HGCAMT and HSGCA, HGCAMT and WHGCAMT, HMM and HSGCA, HMM and WHGCAMT, HSCA and HSGCA and HSCA and WHGCAMT (Figure 5). Furthermore, intra-group hybrids between HGCAMT and HMM, HGCAMT and HSGCA, HMM and HSGCA, HSCA and HGCAMT and HSCA and WHGCAMT were also significant. Similar results were obtained for the economic heterosis of inter- and intra-group crosses of all the methods under *Striga* infestation and optimum environments (Figure S4). However, SCA effects among inter- and intra-group crosses of all the methods under *Striga* infestation and optimum environments were not significant except between intra-group hybrids of HGCAMT and WHGCAMT under *Striga* infestation and HGCAMT and HMM under optimum environment (Figure S5).

Under *Striga* infestation, HSCA, HSGCA, HMM, HGCAMT and WHGCAMT produced 186 and 114, 104 and 196, 192 and 108, 150 and 150 and 100 and 200 total number of inter- and intra-group crosses, respectively. The methods identified 25 and 17, 55 and 154, 74 and 85, 55 and 108 and 59 and 162 number of heterotic and non-heterotic hybrids, respectively. Consequently, those methods had breeding efficiency of 14.2%, 65.7%, 58.6%, 54.3% and 70.0%, respectively (Table 7). Similarly, under optimum environments, 17, 25, 23, 15 and 26 heterotic and 83, 147, 104, 162 and 198 non-heterotic hybrids were identified through HSCA, HSGCA, HMM, HGCAMT and WHGCAMT methods, respectively. The methods indicated breeding efficiency of 48.7, 57.6, 54.1, 52.5 and 62.5, respectively (Table 7).

**Table 7.** Number of heterotic, non-heterotic inter-group, intra-group hybrids and breeding efficiency of HSCA, HSGCA, HMM, HGCAMT and WHGCAMT heterotic grouping methods under *Striga*-infested and optimum environments

| Source | *Striga* infested |  |  |  |  | Optimum |  |  |  |  |
| --- | --- | --- | --- | --- | --- | --- | --- | --- | --- | --- |
|  | HSCA | HSGCA | HMM | HGCAMT | WHGCAMT | HSCA | HSGCA | HMM | HGCAMT | WHGCAMT |
| NHH | 25 | 55 | 74 | 55 | 59 | 17 | 25 | 23 | 15 | 26 |
| NINTGH | 186 | 104 | 192 | 150 | 100 | 207 | 151 | 192 | 126 | 100 |
| NNHH | 17 | 154 | 85 | 108 | 162 | 83 | 147 | 104 | 162 | 198 |
| NINTRGH | 114 | 196 | 108 | 150 | 200 | 93 | 149 | 108 | 174 | 200 |
| BE (%) | 14.2 | 65.7 | 58.6 | 54.3 | 70.0 | 48.7 | 57.6 | 54.1 | 52.5 | 62.5 |

*HSCA: heterotic grouping based on SCA of grain yield; HSGCA: heterotic grouping based on SCA and GCA of grain yield; HMM: heterotic grouping based on molecular marker; HGCAMT: heterotic grouping based on GCA of multiple traits; WHGCAMT: weighted heterotic grouping based on GCA of multiple traits; NHH: number of heterotic hybrids; NINTGH: number of inter-group hybrids; NNHH: number of non-heterotic hybrids; NINTRGH: number of inter-group hybrids; BE: breeding efficiency*

The correlation analysis revealed that grain yield (GYD) was significantly positively correlated with specific combining ability (SCA) effects, mid-parent heterosis (MPH), better-parent heterosis (BPH), and economic heterosis with correlation coefficients of 0.83, 0.99, 0.99 and 1, respectively (Figure 6). Additionally, MPH and BPH were perfectly correlated (r = 1.00***), suggesting they reflect similar genetic contributions to hybrid performance. In contrast, grain yield exhibited significant negative correlations with similarity matrices of WHGCAMT (r = – 0.96***), HGCAMT (r = –0.85***), HSGCA (r = –0.87***), and HMM (r = –0.73*). Furthermore, WHGCAMT was positively associated with HGCAMT (r = 0.74*), HSGCA (r = 0.97***) and HMM (r = 0.55).

**Figure 6.**
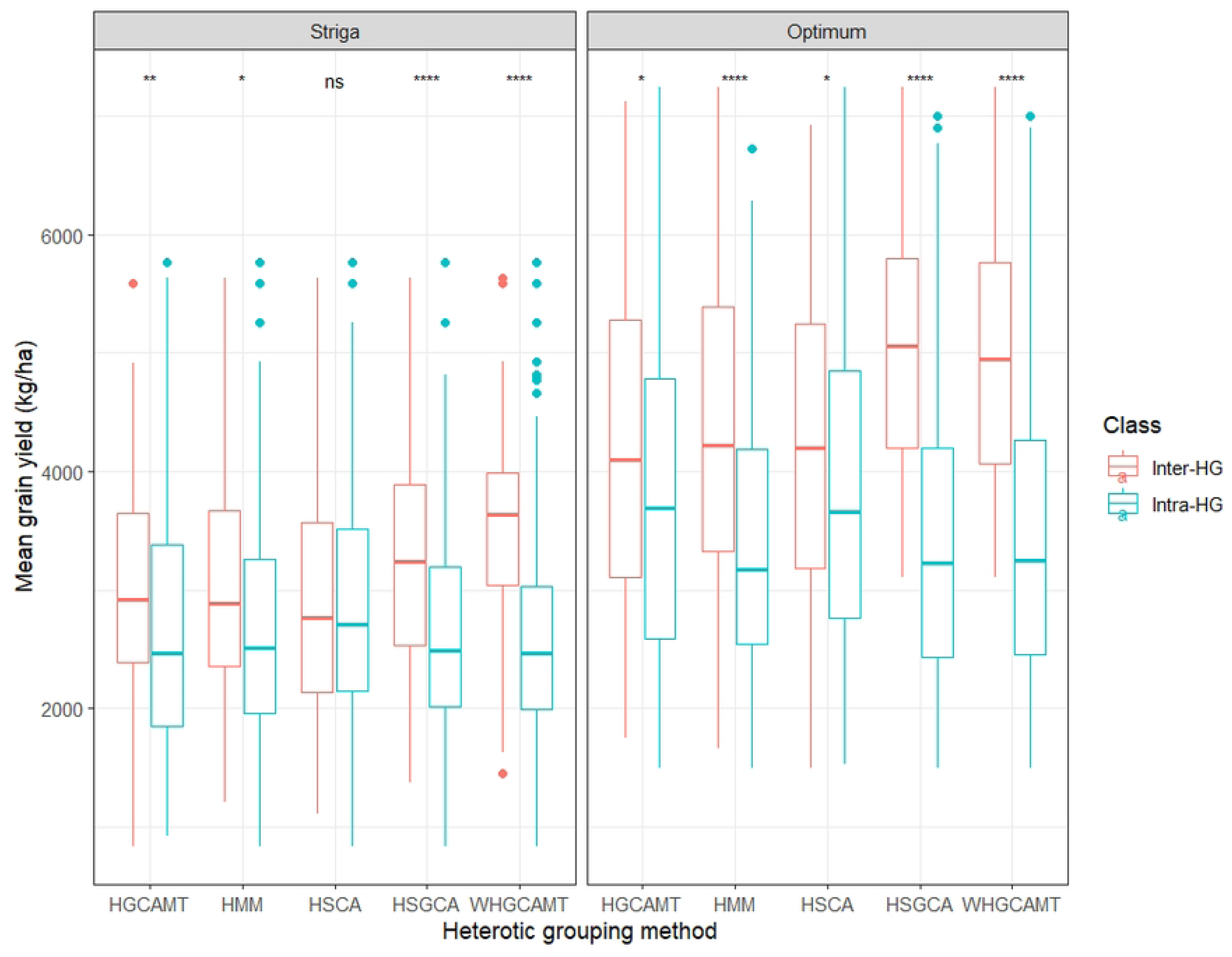
Correlation coefficient for GYD, SCA, MPH, BPH, ECH and the similarity matrices of the 4 different heterotic grouping methods of 300 diallel crosses. *HSGCA: heterotic grouping based on SCA and GCA of grain yield; HMM: heterotic grouping based on molecular marker; HGCAMT: heterotic grouping based on GCA of multiple traits; WHGCAMT: weighted heterotic grouping based on GCA of multiple traits; GYD: grain yield; SCA: specific combining ability effects; ECH: economic heterosis*

## Discussion

In achieving breeding objectives, genetic variability is of prime importance because it provides the basis for selection. Therefore, it is essential to evaluate the genetic variability of the study population. Understanding the extent and nature of variability present in a population helps in choosing appropriate breeding strategies, whether based on additive or non-additive gene action (52,53). Significant genotype mean squares observed for most traits under *Striga* infestation and optimum environments indicated the existence of substantial genetic differences among the study population for improvement. Significant variability observed for the measured traits further provided evidence of inherent genetic differences among the parents and the progenies. This variability can be exploited for improvement of these traits and that the potential of genotypes for selection. The result observed in this study agreed with other findings (21,52,54). The significant mean squares observed for the environment indicated the uniqueness of the test environments to discriminate among the genotypes under *Striga* and optimum environments suggested the need for multi-environment evaluation of the hybrids (55). The significant genotype × environment interaction observed for most measured traits across the test environments revealed genotypic performance varied across environments. This implied that different hybrids should be selected for specific environments, and genotypes that interact less with the environment (i.e., show stability) should be recommended for growth across the test environments (54,56). The significant GCA and SCA mean squares detected for grain yield and most other agronomic traits both under *Striga* infestation and optimum environments indicated that both additive and non-additive gene actions were important in the inheritance of grain yield and other measured traits (32). Variations observed in the GCA mean squares further indicate the availability of genetic variation among the inbreds studied. This indicated the potential for classifying the inbred lines into heterotic groups and identifying lines with high or low favorable alleles that could be useful for population improvement(20) Significant GCA for grain yield and most other traits will facilitate the identification of potentially discriminating inbred testers as well as candidate parental inbreds for traits introgression and hybrid development (32). Significant environment × SCA interaction for grain yield and the majority of the measured traits observed under *Striga* infestation and optimum environments indicated differential performance of the hybrids across environments. This suggests selection of hybrids best suited to a particular environment (57).

Although the non-additive effects were significant, the predominance of additive effects were observed in all the measured traits in this study, where GCA accounted for 95% and 97% of the total genetic variance for grain yield under *Striga*-infested and optimum environments, respectively, indicating that additive gene action was the major determinant of trait expression across test conditions. According to Baker (45) ratios approaching unity suggest high predictability of hybrid performance based on GCA alone. Thus, the inheritance of grain yield and other measured traits was largely governed by additive effects rather than non-additive effects, indicating high breeding value, and highlighting the importance of selecting parents with strong general combining ability. The result agreed with the findings of other authors (10,11).

Moreover, molecular markers were used to assess the level of divergence of the studied population. The observed distribution of the marker across the genome appeared relatively balanced, with SNPs mapped across all 10 chromosomes with chromosome five having the highest number of markers. The PIC of markers averaged 0.40, indicating that the markers used for this study were highly informative and as such capable of providing reliable information on the genotype discrimination, indicating that the SNP markers used will provide sufficient information for heterotic grouping. The SNP markers used in this study were more informative than those of previous studies (58–60). Minor Allele Frequency (MAF) had a mean of 0.28 indicating that a substantial proportion of the inbred population carry rare alleles. The observed heterozygosity (Ho) averaged 0.04 which was very low compared to the expected heterozygosity (He) averaged 0.37. The result further revealed Ho values below 0.01 for more than 95 % of the loci. This indicated that inbred lines used in this study were highly homozygous, indicating the significance of inbreeding and selection effort put in selfing the lines to S_7_. The genetic differentiation observed at the molecular level between (38%) and within (62%) heterotic groups, indicates that while there are clear genetic distinctions of individuals within heterotic groups, a considerable amount of diversity exists between heterotic groups. The differences observed between clusters were higher than previous studies (60–62) indicated that the heterotic grouping based on molecular marker was effective in this study.

The usefulness of inbred lines is attributed to their ability to transmit their characteristics to their progeny. Inbred lines with favorable GCA effects for grain yield and other traits could be use as parents to form a synthetic population that could be improve for stress environments (2). Inbred parents TZEI2248, TZEI2249, TZEI2250, TZEI2251, TZEI18 and TZdEI352 possessed significant positive GCA effects for GYD across *Striga*-infested and optimum environments, indicating that these inbreds could be useful in contributing favorable alleles for improved grain yield under *Striga* and optimum environments. Additionally, those inbred lines possessed significant positive GCA effects for EPP and significant negative GCA effects for SDS, ESP, PASP and EASP, suggesting that they may be useful sources of *Striga* resistance alleles for introgression into susceptible genotypes for genetic enhancement of *Striga* resistance and for the development of superior *Striga*-resistant hybrids and synthetic varieties (32). On the other hand, the inbred lines TZEI2235 and TZEI2238 with significant negative GCA effects for GYD, significant positive GCA effects for SDS, ESP, PASP and EASP are susceptible to *S. hermonthica* and they will contribute unfavorable alleles to their progenies. The observed patterns of GCA effects strengthen the importance of selecting inbred parents that have the ability to transfer both yield potential and *Striga* resistance traits and selection of parents for recycling.

The classification of inbreds into appropriate heterotic groups is fundamental to maximizing their potential as parents in hybrid and synthetic development (7,8,10,15). Four major approaches are currently in use to exploit genetic variability within populations for heterosis and these methods have contributed to the development and commercialization of several outstanding hybrids. However, their efficiency in fully capturing and utilizing genetic variability need to be improved.

The comparative analysis of heterotic grouping methods revealed substantial differences in their effectiveness for classifying inbred lines into heterotic group (Bhat et al., 2024; (Oyetunde et al., 2020; Priscilla et al., 2023). Although, all five methods produced broadly similar grouping trends, but the placement of individual lines varied. A key observation was the higher consistency of the WHGCAMT across contrasting environments. WHGCAMT placed 21 of 25 inbred lines into the same groups under both *Striga* and optimum conditions, compared with 19 for HSGCA, 18 for HGCAMT, and 15 for HSCA. This stability indicates the advantage of incorporating trait weights and accounting for inter-trait correlations, thereby minimizing redundancy and reducing environment-driven reclassification of the inbred lines. The method is flexible, allowing breeders to prioritize traits according to their specific breeding objectives. Besides, the method is scale-independent, accounts for trait correlations thereby enhancing hybrid development. Previous studies have similarly reported that grouping methods based on GCA are generally more stable than those based solely on SCA or molecular markers (10). Contradictory, Bhat et al. (63) reported that the HSCA method was 2.2 and 6.2% more efficient than HSGCA and 10 and 18.1% more efficient than HGCAMT during rainy and post-rainy seasons, respectively. However, their study revealed differential breeding efficiencies of grouping methods and suggested the existence of condition-specific heterotic grouping in maize. WHGCAMT placed all the *Striga-*resistant testers and one susceptible tester in the same heterotic group, while HMM placed all the *Striga*-resistant and susceptible testers in the same group. This indicates that testers possess underlying genetic backgrounds that make them reliable discriminants for grouping (12,65).

A better classification method is one in which heterotic groups allow inter-group crosses to produce more superior hybrids than intra-group crosses (15). It is documented that superior hybrids should arise predominantly from inter-group rather than intra-group crosses (7,15). Although the results showed clear differences between inter- and intra-group hybrids for all the methods except for HSCA, WHGCAMT proved most effective. Under *Striga* infestation, inter-group hybrids identified by WHGCAMT displayed clear advantages in grain yield and economic heterosis compared with intra-group hybrids, followed by HSGCA, HMM and HGCAMT, while HSCA failed to discriminate between the two. This outcome reflects the well-known instability of SCA-based classifications, which are heavily influenced by the interactions between the two parents and hybrid × environment (12,15). Pairwise comparisons of inter- and intra-group hybrids across all methods showed significant differences between WHGCAMT and the other methods in most cases. WHGCAMT consistently produced higher-yielding inter-group and lower-yielding intra-group hybrids. This further underscores the superiority of WHGCAMT over other methods, suggesting that integrating information from multiple traits provides a more reliable classification of inbred lines. This result is consistent with previous findings that multi-trait approaches capture a broader spectrum of genetic variability and better reflect the complex interactions influencing grain yield under stress conditions (2,20,23). The significant differences observed for economic heterosis among the methods provide additional evidence of the utility of WHGCAMT in maximizing heterosis exploitation.

Interestingly, the largely non-significant differences in specific combining ability (SCA) effects, for inter- and intra-group hybrids in most cases, suggest that non-additive effects contribute less to yield expression under both *Striga*-infested and optimum environments. This further strengthens the argument that additive gene effects predominate the expression of the traits, as has been widely reported in maize (12). Thus, WHGCAMT, which emphasizes GCA while accounting for trait correlations and scale-independence, is better aligned with the genetic architecture of hybrid performance.

Breeding efficiency further confirmed the superiority of WHGCAMT, which achieved 70% under *Striga* and 62.5% under optimum conditions, compared to 14.2% and 48.7%, 54.3% and 52.5%, 58.6% and 54.1% and 65.7% and 57.6% for HSCA, HGCAMT, HMM and HSGCA under *Striga* and optimum environments, respectively. These results align with earlier findings that additive-based methods are more predictive of hybrid performance than SCA alone (11,52,65). Marker-based classification (HMM) produced moderate efficiencies, but its poor discrimination under *Striga* stress suggests that molecular distance alone is insufficient, as marker divergence does not always align with functional divergence for stress-related traits (8).

The strong positive correlations observed between grain yield, SCA, and heterosis estimates (MPH, BPH, and ECH) confirm their shared genetic basis and value in predicting hybrid performance. Additionally, the significant negative correlations between grain yield and similarity matrices from WHGCAMT, HGCAMT, HSGCA, and HMM indicate that greater genetic distance between inbred lines tends to translate into superior hybrid performance, strengthening the fundamental idea of inter mating (7,8). Although significant correlations between grain yield and all other methods were detected, WHGCAMT had relatively high value, reinforcing its superiority. The nearly perfect positive correlation between ECH, MPH and BPH further suggests that either of the heterosis may be used interchangeably in evaluating hybrid performance.

## Conclusion

In conclusion, the findings demonstrated a larger proportion of GCA effects of inbreds for grain yield and all other measured traits than those of the SCA effects across the test environments. This suggests that additive gene action played a predominant role than non-additive in the inheritance of the measured traits in the single-cross hybrids evaluated and that GCA was the main component accounting for the differences among the hybrids. Additionally, WHGCAMT offers a more productive and reliable classification for defining heterotic groups among early-maturing maize inbreds, particularly in *Striga*-endemic environments. By weighting key adaptive traits and correcting for correlations among them, one ensures that each trait contributes uniquely context-relevant information to the grouping process. The proposed WHGCAMT heterotic grouping method will enhance the efficiency of hybrid development, ensure greater consistency in inter-group superiority, and ultimately accelerate genetic gain for grain yield and *Striga* resistance in maize breeding programs.

## Data availability statement

The data used/analyzed in this manuscript are publicity available in IITA CKAN repository as follows:

## Genotypic data of the 25 involved in the diallel crosses

https://doi.org/10.25502/bch3-ce59/d

## Phenotypic data of the 300 diallel crosses and 6 checks evaluated in Abuja and Mokwa under Striga-infested and optimum environments

https://doi.org/10.25502/0zkd-zf95/d

## Acknowledgments

The authors are grateful to the IITA Maize Improvement Programme and Bioscience staff for providing technical assistance.

## Supplementary Information

Table S1. Description of the 25 inbred lines used in this study

Table S2. SCA effects of the crosses for grain yield and other Striga-adaptive traits evaluated under Striga-infested environment in Abuja and Mokwa in Nigeria

Table S3. SCA effects of the crosses for grain yield and other agronomic traits evaluated under optimum environment in Abuja and Mokwa in Nigeria

Table S4. Mean grain yield and other agronomic traits of the inbred parents evaluated under Striga infestation in Abuja and Mokwa in Nigeria

Table S5. Mean grain yield and other Striga-adaptive traits of the hybrids evaluated under Striga infestation in Abuja and Mokwa in Nigeria

Table S6. Mean grain yield and other agronomic traits of the hybrids evaluated under optimum environment in Abuja and Mokwa in Nigeria

Table S7. Heterotic grouping of the inbred parents based on five different methods under Striga infestation and optimum environments

Figure S1a. Silhouette K-means analysis showing optimum number of clusters and the dendrogram of 25 maize inbred lines constructed from GCA effects of multiple traits (HGCAMT) using Ward’s minimum variance cluster analysis.

Figure S1b. Silhouette K-means analysis showing optimum number of clusters and the dendrogram of 25 maize inbred lines constructed from weighted GCA effects of multiple traits (WHGCAMT) using Ward’s minimum variance cluster analysis

Figure S1c. Silhouette K-means analysis showing optimum number of clusters and the dendrogram of 25 maize inbred lines constructed from 1696 SNP markers using Ward’s minimum variance cluster analysis

Figure S1d. Silhouette K-means analysis showing optimum number of clusters and the dendrogram of 25 maize inbred lines constructed from GCA effects of multiple traits (HSGCA) using Ward’s minimum variance cluster analysis

Figure S2. Pair comparison between intra- and inter-group crosses for economic heterosis of five different heterotic grouping methods under Striga-infested and optimum environments

Figure S3. Pair comparison between intra- and inter-group crosses for SCA effects of five different heterotic grouping methods under Striga-infested and optimum environments

Figure S4. Pair comparison among the heterotic grouping methods for economic heterosis produced by intra- and inter-group crosses under Striga-infested and optimum environments

Figure S5: Pair comparison among the heterotic grouping methods for SCA effects produced by intra- and inter-group crosses under Striga-infested and optimum environments

